# Characterization and Mitigation of a Simultaneous Multi-Slice fMRI Artifact: Multiband Artifact Regression in Simultaneous Slices

**DOI:** 10.1101/2023.12.25.573210

**Authors:** Philip N. Tubiolo, John C. Williams, Jared X. Van Snellenberg

**Affiliations:** Department of Biomedical Engineering, Stony Brook University, Stony Brook, NY 11794; Department of Psychiatry and Behavioral Health, Renaissance School of Medicine at Stony Brook University, Stony Brook, NY 11794; Department of Psychology, Stony Brook University, Stony Brook, NY 11794

## Abstract

Simultaneous multi-slice (multiband) acceleration in fMRI has become widespread, but may be affected by novel forms of signal artifact. Here, we demonstrate a previously unreported artifact manifesting as a shared signal between simultaneously acquired slices in all resting-state and task-based multiband fMRI datasets we investigated, including publicly available consortium data from the Human Connectome Project (HCP) and Adolescent Brain Cognitive Development (ABCD) Study. We propose Multiband Artifact Regression in Simultaneous Slices (MARSS), a regression-based detection and correction technique that successfully mitigates this shared signal in unprocessed data. We demonstrate that the signal isolated by MARSS correction is likely non-neural, appearing stronger in neurovasculature than grey matter. Additionally, we evaluate MARSS both against and in tandem with sICA+FIX denoising, which is implemented in HCP resting-state data, to show that MARSS mitigates residual artifact signal that is not modeled by sICA+FIX. MARSS correction leads to study-wide increases in signal-to-noise ratio, decreases in cortical coefficient of variation, and mitigation of systematic artefactual spatial patterns in participant-level task betas. Finally, MARSS correction has substantive effects on second-level t-statistics in analyses of task-evoked activation. We recommend that investigators apply MARSS to multiband fMRI datasets with moderate or higher acceleration factors, in combination with established denoising methods.

## 1. Introduction

### 1.1 Principles and Advantages of Multiband fMRI

Simultaneous multi-slice (multiband; MB) fMRI has become a prevalent fMRI acquisition acceleration technique that partially mitigates the limitations posed by standard single-slice (single-band) EPI acquisition^1,2,3^. In MB fMRI acquisition, composite radiofrequency pulses comprising central frequencies (with bandwidths that determine slice thickness) that match the precession frequency of hydrogen nuclei in several axial sections along the field of view are transmitted to acquire signal from several slices in a single readout; the number of slices acquired simultaneously is denoted by the MB acceleration factor. During readout, simultaneously acquired slices are “stacked” on top of each other and phase-shifted along the phase-encoding direction to reduce the amount of overlapping signal between slices^4,5^. Slices are then unmixed using matrix inversion techniques that utilize radiofrequency coil sensitivity maps acquired prior to image acquisition. Overall, MB acceleration allows for substantive increases in spatiotemporal resolution that reduce signal-to-noise ratio but improve the spatial precision of neural signal estimates^1^. These improvements in data quality have led to its widespread adoption in both individual research laboratories and in large consortiums that have produced publicly available datasets, such as the Human Connectome Project (HCP)^6,7^, UK Biobank study^8^, and Adolescent Brain Cognitive Development (ABCD) Study^9^.

### 1.2 Previously Discovered Multiband-Specific Artifacts and Mitigation Methods

As MB acceleration has gained adoption, several sources of contamination stemming from MB acquisition have been identified. Todd and associates identified a source of shared signal among voxels in simultaneously acquired slices that were overlapped during readout, coined “Interslice leakage”^10^. It was determined that this artifact could be substantially reduced by using Split-Slice Generalized Autocalibrating Partial Parallel Acquisition (GRAPPA)^11^ image reconstruction over other techniques, such as Slice-GRAPPA^12^. McNabb and associates subsequently identified that interslice leakage artifacts are exacerbated by eye movement, even when Split-splice GRAPPA reconstruction is used, by having participants forcefully blink during resting-state MB fMRI acquisition^13^.

Additionally, MB fMRI acquisition poses unique challenges in the estimation and removal of artifacts due to in-scanner participant motion, as the higher sampling rate enabled by these sequences facilitates the capture of additional nuisance signals^14,15,16,17,18,19,20,21^, including respiratory motion and pseudomotion, or factitious head motion observed in the phase-encode direction as a consequence of lung expansion distorting the B_0_ field^16^.

### 1.3 Discovery of a Shared Artifact Signal in Simultaneously Acquired Slices

In working with resting-state and task-based MB fMRI data in our own laboratory, we observed periodic banding patterns in carpet plots^22^ that display mean slice-wise timeseries. In investigating the source of this banding pattern, we discovered that raw, unprocessed MB fMRI data display elevated Pearson correlations between the average timeseries in simultaneously acquired slices, visible as diagonal bands in a correlation matrix of average slices signals in the original scanner space (**Figure 1**). We speculated that this indicated a shared signal source between simultaneously acquired slices that warranted further investigation, given that there is no biological basis for true neural signal fluctuations to be shared between arbitrarily located, spatially disparate, axial slices over and above that which is shared with other slices in the image. Therefore, we believe that these elevated correlations represent a BOLD signal artifact: a non-neural, structured signal source that is uniquely detectable in MB accelerated fMRI. Notably, our use of the term “artifact” to refer to this signal may differ somewhat from some definitions of an “image artifact” or “multiband artifact.” We do not mean to assert that this artifact is somehow produced or created by the image sequence or multiband implementation, although this remains a possibility. Rather, we use the term “multiband artifact” to describe an undesirable, non-neural, signal source that negatively impacts data quality and that is present in BOLD multiband imaging. Indeed, this signal source could still exist in singleband fMRI, but if so it would be undetectable because no two slices are acquired simultaneously.

**Figure 1.**
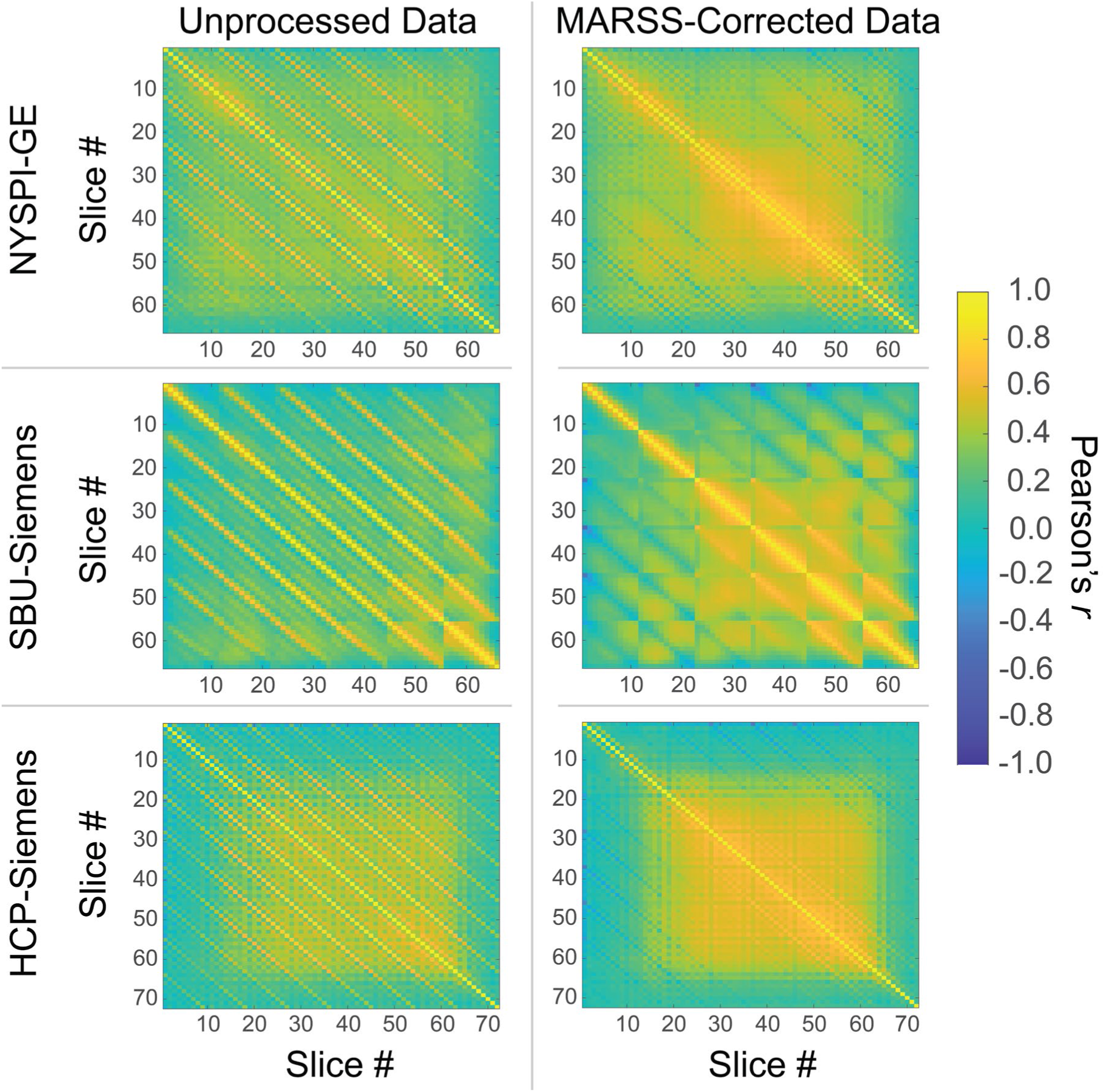
Pearson correlation matrices between average signal in all slice pairs in unprocessed and MARSS-corrected task-based data from NYSPI-GE, SBU-Siemens, and HCP-Siemens.

Consequently, we detail the identification of this novel artifact signal across several resting-state and task-based MB fMRI datasets, including data from two large, publicly available, datasets^6,7,9^, and three major scanner manufacturers: Siemens (Munich, Germany), General Electric (GE; Boston, MA), and Philips (Andover, MA). We propose a regression-based technique to reduce the presence of the shared signal between simultaneously acquired slices, via estimation and removal of the shared signal in unprocessed MB fMRI data files, which we term Multiband Artifact Regression in Simultaneous Slices (MARSS). This estimation method is based on the idea that, provided major sources of shared signal (i.e., global signal and motion-related signal) are separately accounted for, averaging together a sufficient number of voxels containing the same artifact signal (i.e., all voxels from all simultaneously acquired slices) will converge on the artifact signal itself, as true neural signals from disparate brain regions will “average out.”

Furthermore, we determine whether this artifact signal is distinct from previously identified interslice leakage artifacts. We also investigate the spatial, temporal, and spectral properties of the isolated artifact signal. We then explore whether denoising via spatial Independent Components Analysis with FMRIB’s ICA-based Xnoiseifier (sICA+FIX)^23,24^ mitigates this artifact signal when used both in isolation and in tandem with MARSS. Finally, we perform a working memory task analysis in both processed and unprocessed MB fMRI to highlight the effects of MARSS correction on spatial patterns of estimates of task-evoked activation. We predicted that the artifact signal may originate from a combination of fluid motion and periodic mechanical noise during acquisition. Further, we hypothesized that noise reduction due to artifact removal would result in increased magnitude of spatial t-statistics in between-participants analyses of task-based fMRI, most likely as a result of reduced variance due to removal of spurious artifact signals.

## 2. Materials and methods

### 2.1 Functional Magnetic Resonance Imaging Datasets

To comprehensively evaluate the presence of the artifact signal across a range of scanner platforms and scan parameters, we utilized phantom and *in vivo* MB fMRI data from several sources, including samples from two publicly available datasets (HCP and ABCD), and two in-house samples from the New York State Psychiatric Institute (NYSPI) and Stony Brook University (SBU). A comprehensive data summary including scan acquisition parameters can be found in **Table 1**, and demographic information for each dataset in **Table S1**.

**Table 1.**
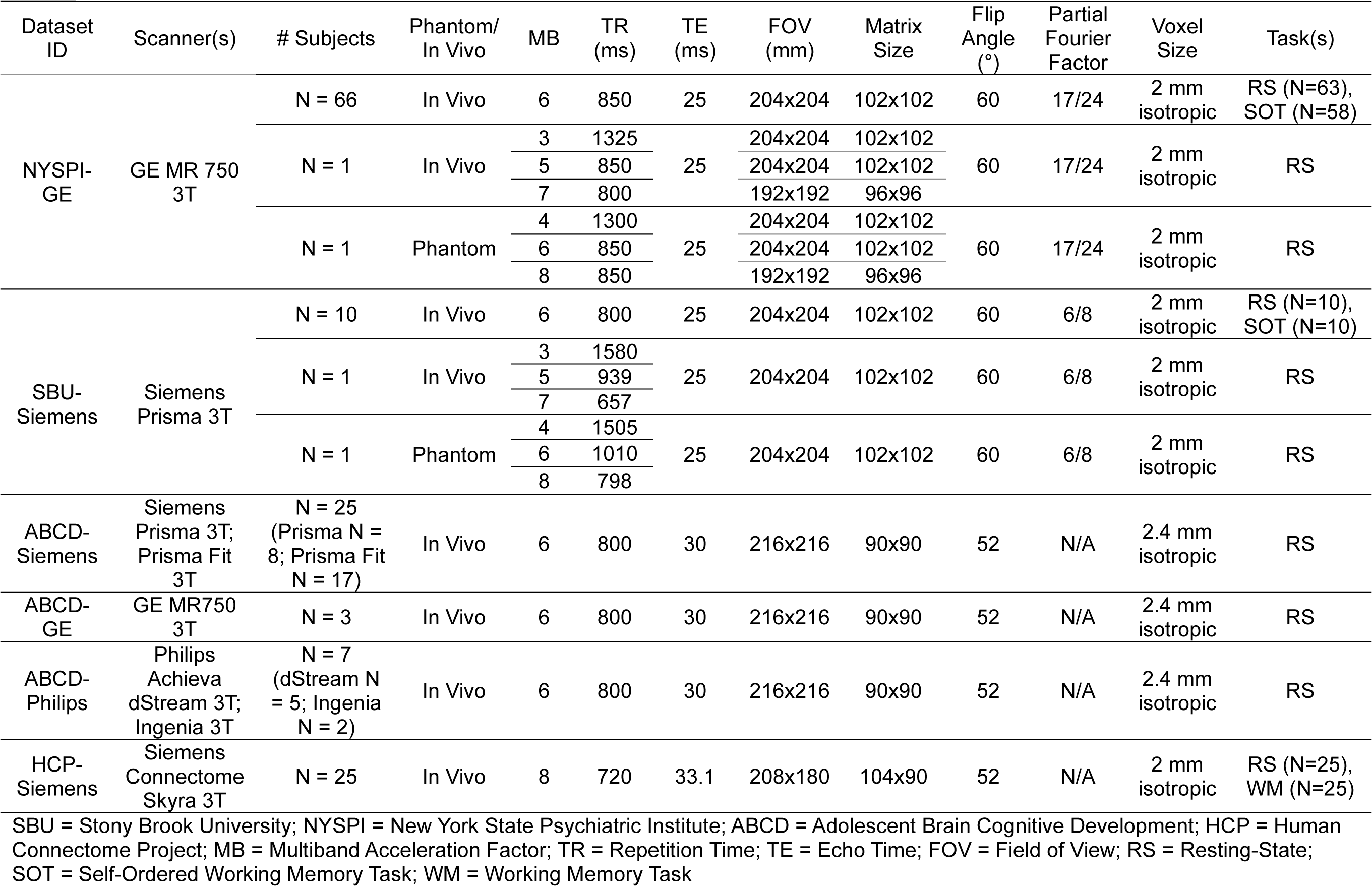
Datasets used in this study. Datasets are referred to by their Dataset ID throughout the manuscript.

All procedures described in this study that involve the acquisition, curation, and analysis of data from human participants at NYSPI and SBU were approved by the Institutional Review Boards of NYSPI -Columbia University Department of Psychiatry and SBU, respectively. All participants provided written consent prior to their participation in study procedures.

#### 2.1.1 Resting-State *in Vivo* Datasets

We first employed unprocessed resting-state MB fMRI data acquired at NYSPI (NYSPI-GE; n = 63), with between 1 and 4 runs per participant (1 participant had 1 run, 4 had 2 runs, 9 had 3 runs, and 49 had 4 runs). Another resting-state dataset was acquired at SBU (SBU-Siemens; n = 10), with 4 or 6 runs per participant (1 participant had 6 runs). NYSPI-GE data was collected using a Nova Medical (Wilmington, MA) 32-channel head coil, while SBU-Siemens data used a Siemens 64-channel head-and-neck coil.

We also used resting-state data from a subset of the HCP (HCP-Siemens)^6,7^ 500 Subjects Release (n = 25) and the ABCD Study (ABCD-Siemens, ABCD-GE, and ABCD-Philips; n = 35)^9^. All participants in the HCP-Siemens dataset had 4 runs of resting-state data, while participants in the ABCD datasets had a minimum of 1 and maximum of 5 runs of resting-state data.

To evaluate the relationship between artifact presence and MB factor *in vivo*, one participant each from SBU-Siemens and NYSPI-GE underwent 1 resting-state run for each MB factor of 3, 5, and 7.

#### 2.1.2 Task-Based Datasets

Up to 4 runs of task-based MB fMRI data were acquired from participants in the SBU-Siemens dataset (n = 10; 1 participant had 3 runs) and a subset of the NYSPI-GE dataset (n = 58; 8 participants had 1 run, 4 had 2 runs, 14 had 3 runs, and 32 had 4 runs) using identical acquisition parameters to the resting-state data. Participants performed the self-ordered working memory task^25^ (SOT; described in the Task Procedures section of **Supplementary Methods**). Additionally, each HCP-Siemens participant (n = 25) performed 2 runs of working memory task-based data^26^ with identical acquisition parameters to HCP-Siemens resting-state data.

#### 2.1.3 Phantom Datasets

Phantom data was acquired to investigate both the non-neural origin of the artifact signal and the relationship between artifact presence and MB factor. Three runs of a Siemens D165 Spherical Phantom containing a Nickel sulfate solution were acquired at SBU, and three runs of an agar-filled Functional Biomedical Informatics Research Network (FBIRN) spherical stability phantom^27^ were acquired at NYSPI.

### 2.2 Identification of Artifact Signal in Simultaneously Acquired Slices

A shared artifact signal across simultaneously acquired slices was identified separately in each run of MB fMRI data across all datasets. For each run, the average timeseries in each axial slice was obtained. To remove the impact of participant head motion on raw slice timeseries data, each slice timeseries was then taken to be the residuals of a multiple linear regression modeling the average slice timeseries as a function of six rigid-body head motion parameters, their squares, derivatives, and squared derivatives. The presence of a shared artifact signal across simultaneously acquired slices was quantified as the difference in the mean Pearson correlation between the timeseries in simultaneously acquired slices and the mean correlation between the pairs of slices immediately adjacent to the each slice in the simultaneous slice set (hereon simply referred to as “adjacent-to-simultaneous” slices), averaged across all slices in a volume. All Pearson correlations are Fisher Z-transformed prior to averaging or subtracting and transformed back to Pearson *r* before being reported throughout this manuscript. This procedure was performed before and after artifact correction techniques were used to quantify the magnitude of artifact and success of artifact removal. Statistical significance of elevated correlations between simultaneously acquired slices in uncorrected data and the reduction in these correlations in MARSS-corrected data were determined via Wilcoxon signed-rank tests^28,29^.

### 2.3 Multiband Artifact Regression in Simultaneous Slices (MARSS)

#### 2.3.1 MARSS Estimation of Artifact Signal

We developed a regression-based method, MARSS, to estimate the magnitude of, and remove, the artifact signal shared across simultaneously acquired slices for each run of MB fMRI data. First, given a run with *N*_*s*_ slices and a MB acceleration factor *MB_f_*, the number of simultaneously acquired slices is given by:

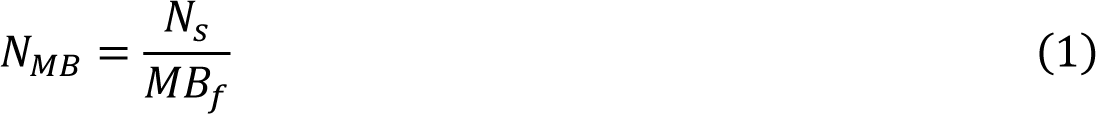

Given a slice *j* from this run, we define the set of slices acquired simultaneous to *j* as:

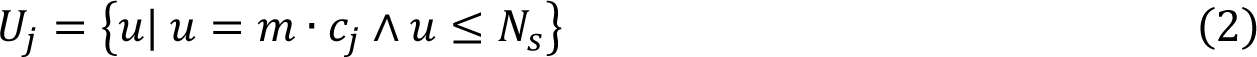

where *N*_*s*_ is the total number of slices in the volume, *m* ∈ ℕ^+^, and:

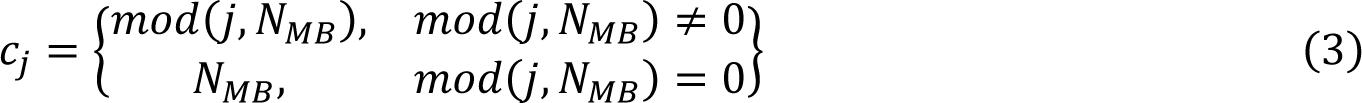

Next, we define the average timeseries across all voxels *d*_*i*,*u*_ in slices simultaneously acquired to slice *j*, but excluding slice *j*, as:

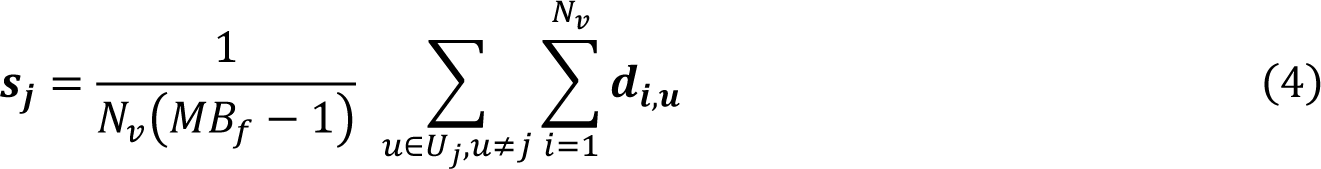

where *N*_*v*_ is the number of voxels in each slice. We also define the average global signal for all slices that were not acquired simultaneously to slice *j* as:

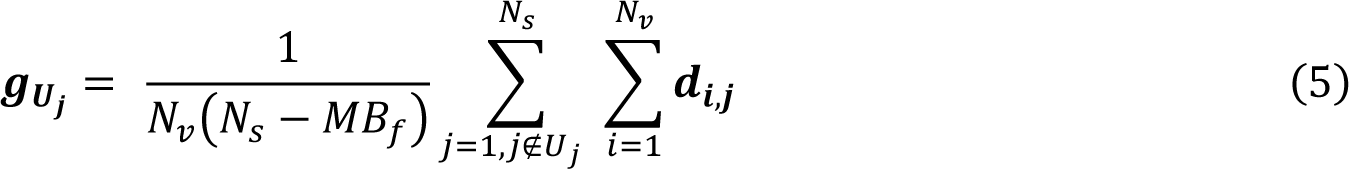

To estimate the artifact signal via regression, we construct a design matrix:

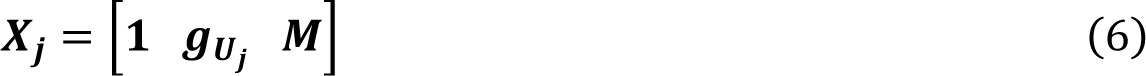

where **1** is a vector of 1s with length equal to the run timeseries (i.e., number of volumes), and ***M*** is a matrix of nuisance parameters, including six rigid-body head motion parameters and their squares, derivatives, and squared derivatives. Next, we regress ***X***_***j***_ onto ***s***_***j***_:

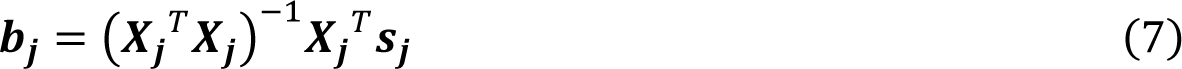

and estimate the artifact timeseries for slice *j* as the residuals of that regression, or:

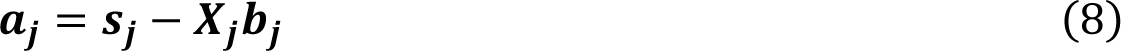

#### 2.3.2 MARSS Removal of Artifact Signal

Given ***a***_***j***_, the artifact signal estimate for slice *j*, we construct a design matrix to estimate the artifact signal contribution to each voxel *i* in slice *j*:

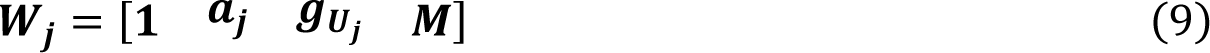

and regress it onto the voxel-wise timeseries ***d***_***i***,***j***_:

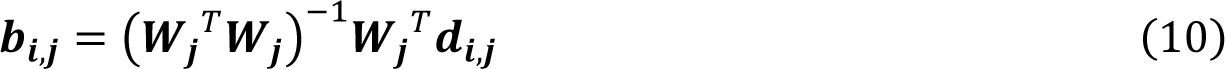

The voxel-wise artifact signal estimate ***a***_***i***,***j***_ can be defined as the product of the columns of ***b***_***i***,***j***_ and ***W***_***j***_ that correspond to the artifact estimate, ***a***_***j***_ (the second column):

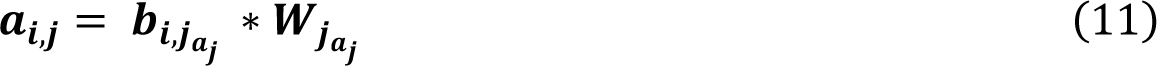

Lastly, we obtain the corrected timeseries ***d***^′^ of voxel *i* in slice *j* as:

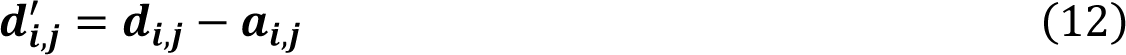

Following MARSS correction, a new fMRI timeseries was saved as a NIFTI image, using the ***d***^′^_*i,j*_ timeseries at each voxel in place of the original data. In addition, the spatial distribution of the artifact estimated by MARSS, ***A***, was determined by taking the mean (over timepoints, *t*) of the absolute value of the artifact signal at each voxel:

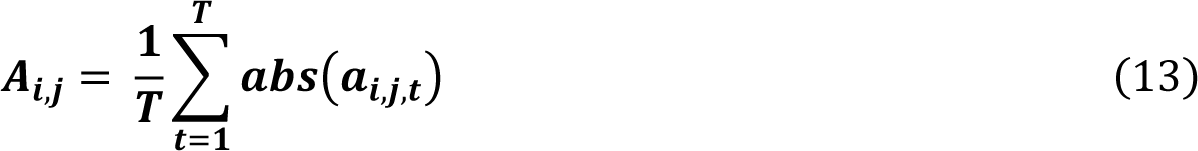

#### 2.3.3 Alternative Artifact Estimation and Removal Methods

In a modification to the technique described above, artifact signal was estimated using only background signal. This was performed by masking out the brain prior to artifact estimation using whole-brain masks generated by SynthStrip^30^ included in FreeSurfer version 7.3.0^31^.

Masks were dilated by one voxel prior to use. MARSS correction and removal remained otherwise unchanged in this modification.

### 2.4 Characterization of Isolated Artifact Signal

Following MARSS artifact estimation and removal, we sought to investigate the spectral, temporal, and spatial properties of the isolated artifact signal to provide evidence for its potential source.

#### 2.4.1 Frequency Spectral Analysis

For each run in a given dataset, the average, whole-image, isolated artifact timeseries was mean-centered. All runs were then concatenated to form a single vector, with zero-padding of the length of the timeseries included after each sample. Welch’s power spectral density^32^ (PSD) was calculated on the concatenated timeseries with a Hanning window applied to each individual timeseries, and 50% overlap between samples. To quantify the average relationship between PSD and frequency in the roughly linear portion of the spectrum (with a lower frequency bound of -2 log[Hz]), a linear regression was performed on log-log transformed PSD that had been weighted by the inverse kernel density of the spectrum. The slope of this regression is useful in estimating colored noise characteristics of the artifact signal.

#### 2.4.2 Autocorrelation

To investigate the potential periodicity of the isolated artifact signal, we calculated its autocorrelation as a function of lag, with each lag representing a temporal shift in the signal of one TR. In each dataset, the isolated artifact timeseries, obtained from Equation 11, above, was spatially averaged to produce a single mean artifact timeseries. This timeseries was then mean-centered and linearly detrended before calculating the autocorrelation up to 20 lags (or a shift of 20 TRs) to sufficiently account for the period of physiological signals that may be components of the artifact signal. Autocorrelations were then averaged across runs in each dataset.

#### 2.4.3 Associations between Artifact Magnitude and In-Scanner Motion

We sought to determine whether the artifact magnitude, measured as the elevation in Pearson correlation in simultaneously acquired slices compared to adjacent-to-simultaneous slices, was associated with measures of in-scanner subject motion. These associations were assessed in resting-state and task-based data from the NYSPI-GE dataset. For each subject, the mean artifact magnitude was calculated as the average of the Fisher Z-transformed simultaneously acquired slice correlation elevations across runs. In-scanner motion was quantified either by 1) framewise displacement (FD), calculated as the sum of the absolute values of the derivative of six motion parameters (X, Y, Z, roll, pitch, and yaw), and 2) DV, calculated as the root-mean-square of the derivative of the timeseries across all in-brain voxels^22^. The mean and median of each motion measure were calculated across runs for each subject, and separate robust linear regressions were used to predict artifact magnitude from each mean and median motion measure.

#### 2.4.4 Interslice Leakage Artifact Analysis

Interslice leakage artifacts have been previously shown to contaminate estimates of neural signal in MB fMRI^13^. They arise as a consequence of the imperfect “unmixing” of signals in simultaneously acquired slices that are originally stacked and phase-shifted by a set fraction of the field of view (FOV) immediately preceding reconstruction. Interslice leakage artifacts, however, typically comprise a shared signal between two voxels that were overlaid during acquisition, rather than entire slices. We sought to demonstrate that the artifact detected and removed by MARSS is not a product of interslice leakage.

In resting-state MB fMRI data in the SBU-Siemens dataset, it was known that simultaneously acquired slices were phase-shifted by multiples of 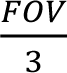 voxels (that is, multiples of 34 voxels) relative to each other prior to reconstruction. For each voxel, the contribution to its signal due to the voxels corresponding to the phase shift was estimated and removed via regression; that is, each voxel in the image was subjected to a regression in which the signals in all voxels that were “stacked” on top of that voxel were regressed out. Although this is destructive to the true underlying neural signal in each voxel, it was employed as a method to determine whether the MARSS artifact remained following removal of slice leakage effects from the data. The presence of the shared artifact in simultaneously acquired slices that remained after removing any potential interslice leakage was then determined using prior methods (see above, *2.2 Identification of Artifact Signal in Simultaneously Acquired Slices*). It is important to note that this procedure is not intended to be a step of MARSS, but rather a separate procedure to demonstrate that the MARSS-corrected artifact signal is independent of interslice leakage.

Additionally, we calculated the Pearson correlation between timeseries in voxels of simultaneous slices that were expected to overlap due to the phase shift during image reconstruction. This was done prior to and following MARSS correction, as well as in data that had undergone slice leakage regression but not MARSS correction, to further demonstrate that MARSS is removing artifact signal independent of interslice leakage.

#### 2.4.5 Effect of Slice Timing Correction on Artifact Magnitude

In order to assess the effect of performing slice timing correction (STC) on artifact presence and magnitude, we performed STC on the unprocessed, resting-state, SBU-Siemens data using SPM12^33,34^. Artifact presence was then evaluated using prior methods (see above, *2.2 Identification of Artifact Signal in Simultaneously Acquired Slices*).

#### 2.4.6 Analysis of Artifact Spatial Distribution

We assessed the average spatial distribution of the isolated artifact signal in resting-state data in NYSPI-GE and SBU-Siemens. For each run, the MARSS estimate of the 3D spatial distribution of artifact was obtained using Equation 13, above. Each run’s spatial distribution was warped and resampled into MNI152Nlin6Asym template space^35^ by the single-step resampling procedure within the HCP Minimal Preprocessing Pipeline (OneStepResampling.sh)^36^. This was performed after fMRI preprocessing and utilized previously calculated gradient distortion corrections and affine transformations calculated during that process (see below, *2.6.3 fMRI Preprocessing*). To determine how the magnitude of signal fluctuations due to artifact was impacted by the underlying signal intensity (e.g., brain matter versus empty space), this process was repeated after rescaling each unprocessed average artifact distribution to reflect the percentage of artifact signal relative to mean signal intensity on a per-voxel basis.

#### 2.4.7 Temporal Signal-to-Noise Ratio

Following artifact removal, we assessed the resulting change in whole-brain temporal signal to noise ratio (tSNR) in both unprocessed and preprocessed resting-state and task-based data. For each run, tSNR in voxel *i* of slice *j* was calculated as:

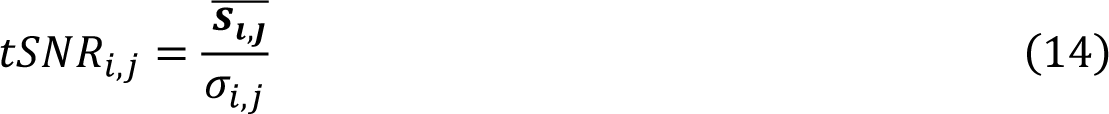

where 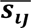 is the mean signal and *σ*_*ij*_ is the standard deviation of the signal^37,38^. In unprocessed data, voxel-wise tSNR estimates were averaged within dilated, whole-brain masks generated for each run by SynthStrip^30^ and compared between corrected and uncorrected data. In preprocessed data, mean tSNR difference and percent difference maps were generated separately for resting-state and task-based data in NYSPI-GE and SBU-Siemens. Significance of the elevation in tSNR across each dataset was determined via Wilcoxon signed-rank tests.

### 2.5 Evaluation of Denoising via sICA+FIX

Since we do not intend for MARSS to be a standalone denoising solution for multiband fMRI, we investigated its efficacy alongside sICA+FIX denoising^23,24^, which is currently implemented in all publicly available, HCP resting-state data^6,7,36,39^. We were particularly interested in the magnitude of the observed artifact in simultaneously acquired slices remaining in the data following sICA+FIX denoising.

#### 2.5.1 Implementation of sICA+FIX in Unprocessed, Resting-State Data

For each resting-state run in the HCP-Siemens dataset, unprocessed fMRI data and associated motion parameters, derivatives, squares, and squared derivatives were first linearly detrended. Independent component (IC) regressors were taken from the previously calculated mixing matrix (*melodic_mix*) released with the *Resting State fMRI 1 FIX-Denoised (Extended)* and *Resting State fMRI 2 FIX-Denoised (Extended)* packages from ConnectomeDB^39^. IC regressors were then z-scored, and the 24 detrended motion parameters were regressed out of both the mixing matrix and the detrended fMRI data. FIX-cleaned data was obtained by regressing all IC regressors onto the fMRI data and calculating the residuals with respect to noise components, only. This procedure was performed in both uncorrected, unprocessed data and MARSS-corrected data.

#### 2.5.2 Subspace Decomposition of sICA+FIX Denoised Data

Denoising via sICA+FIX estimates and removes structured noise components but does not model unstructured or random noise components. These unmodeled components will contain mostly Gaussian noise, but may also contain some weakly structured signals that were not successfully modeled by sICA^15^. Therefore, the denoised data can be decomposed to a sum of a “neural signal” subspace and an unstructured, “random noise” subspace. We performed this decomposition following methods described elsewhere (see Supplementary Materials in^15^) to determine which data subspace contains artifactually elevated correlations between simultaneously acquired slices that are corrected by MARSS. The random noise subspace was obtained by residualizing uncorrected data with respect to all IC regressors, regardless of their classification. The neural signal subspace was then obtained by subtracting the random noise subspace from the sICA+FIX denoised data (i.e., data in which IC regressors classified as noise components are removed). In addition to this decomposition, the signal estimated by sICA+FIX noise components was also evaluated for simultaneous slice artifact presence. Finally, global signal regression (GSR) was performed on the neural subspace to determine whether the residual artifact in this subspace could be captured as global structured noise or signal^15^. Global signal was estimated as the mean timeseries across all in-brain voxels, as determined by previously generated brain masks (see above, *2.3.3 Alternative Artifact Estimation and Removal Methods*).

In each subspace, artifact magnitude was assessed using methods described previously (see above, *2.2 Identification of Artifact Signal in Simultaneously Acquired Slices*), with the only modification being that correlations were only calculated between slices in which all runs of all participants contained in-brain voxels. Correlation matrices of the difference between uncorrected and MARSS-corrected subspaces were also calculated to determine whether structured correlations beyond those in simultaneously acquired slices, specifically those that could reflect neural signal, were being removed by MARSS.

Following the calculation of data subspaces, we performed a variance decomposition of the raw data prior to and following MARSS correction, which is described in detail in the **Supplementary Methods**.

### 2.6 Task-Based Analysis

We sought to determine the effects of MB artifact correction on the spatial distribution of statistical maps resulting from task-based modeling (see **Supplementary Methods**). We first performed three within-participants analyses on unprocessed, working memory task-based MB fMRI data in NYSPI-GE (n=58), SBU-Siemens (n=10), and HCP-Siemens (n = 25) before and after MARSS correction. We then performed separate between-participants analyses on preprocessed SBU-Siemens and NYSPI-GE data to highlight the effects of artifact correction on cortical, group-level t-statistics.

#### 2.6.1 Within-Participants Task Modeling

Prior to modeling, each fMRI run was normalized to reflect percent signal change on a per-voxel basis, ***n_i,j_*** = 100 * ***s_i,j_/s_i,j̅_***, where *s_i,j_* is the timeseries and ***s_ij̅_*** is the mean signal in voxel *i* in slice *j*. Each task was modeled using a single-parameter canonical hemodynamic response function (HRF) in SPM12^33,34^. For the SOT, regressors of interest were those reflecting correct trials in the control task or each of the first seven steps of the SOT, modeled separately. For the visual n-back task, a block design was utilized to model all 0-back and 2-back conditions. Trial timings were extracted from event files (“EVs”) distributed with HCP task data. Nuisance parameters included six motion parameters, their squares, derivatives, and squared derivatives, as well as spike regressors to remove high motion volumes, without being convolved with the HRF. Spike regressors were identified using run-adaptive, generalized extreme value, low-pass filtered DVARS (GEV-DV) thresholds and chosen to identify approximately 3% of total volumes across each dataset^18^. The GEV-DV parameters used for each dataset were 28 in NYSPI-GE, 21 in SBU-Siemens, and 42 in HCP-Siemens.

Following model estimation, whole-brain mean squared residuals were obtained for each participant as a measure of within-participant model error and compared in uncorrected and MARSS-corrected data via Wilcoxon signed-rank test.

#### 2.6.2 Spatial Frequency Analysis of Unprocessed Task fMRI

We performed a spatial frequency analysis along the slice direction in each task dataset to evaluate the presence of periodic patterns of task betas corresponding to simultaneous slice spacing both before and after artifact removal. This was performed separately for task and control conditions in each task. One participant in SBU-Siemens and 7 participants in NYSPI-GE were excluded from this analysis (and subsequent analyses of preprocessed data; see below, *2.5.4 Between-Participants Analysis*) for not sufficiently performing all task conditions.

Betas for each condition were averaged across the x-and y-dimensions, resulting in a single value per slice, and mean-centered. Frequency spectral analysis was then performed using previously described methods (see above, *2.4.1 Frequency Spectral Analysis*).

#### 2.6.3 fMRI Preprocessing

Corrected and uncorrected data in SBU-Siemens and NYSPI-GE were preprocessed using the HCP Minimal Preprocessing Pipeline (MPP)^6,36^ Version 4.2.0. Briefly, data underwent processing through the *PreFreeSurfer, FreeSurfer, PostFreeSurfer, fMRIVolume,* and *fMRISurface* pipelines. A critical step in the HCP preprocessing is its ribbon-constrained volume to surface mapping algorithm^36^. Notably, this step chooses to exclude voxels with locally high coefficients of variation (COV) from the mapping process. We quantified the mean, voxel-wise change and percent change in COV before and after MARSS correction within the cortical ribbon isolated for each participant during anatomical volume preprocessing to determine whether MARSS increases the probability that cortical voxels will be included in volume to surface mapping.

Using the preprocessed outputs in CIFTI format (*dtseries.nii*), cortical surface GIFTI files and subcortical NIFTI files were generated, smoothed with a 4 mm full width at half maximum Gaussian kernel, and normalized to reflect percent signal change on a per-greyordinate basis.

#### 2.6.4 Between-Participants Analysis

First level modeling was performed for the SOT as previously described (see above, *2.5.1 Within-Participants Task Modeling*), with the exception of using cortical surfaces and subcortical volumes rather than pure volumetric data. Task contrasts for each participant were calculated as the average beta estimate from the first seven steps of the SOT minus the control condition.

Contrasts from corrected and uncorrected data were analyzed separately in between-participants analyses using Permutation Analysis of Linear Models (PALM)^40,41,42,43^ alpha119. Analyses in PALM were performed using 1024 and 20000 sign flips in SBU-Siemens and NYSPI-GE, respectively, both with and without threshold-free cluster enhancement (TFCE)^44^. T-statistic result images with and without TFCE were subtracted to obtain differences in t-statistics resulting from artifact removal. This subtraction was also repeated after taking the absolute value of t-statistics in corrected and uncorrected data where uncorrected data had negative t-statistics, yielding difference maps reflecting the magnitude of the change in t-statistics (i.e., so that a decrease in signal in an area showing deactivation would appear as an *increase* in t-statistic magnitude), rather just the direction of the change.

In addition to this, we quantified the change in significant greyordinates resulting from MARSS correction. T-statistic maps were thresholded at p < 0.05, Šidák-corrected^45^, and the number of significant greyordinates prior to and following MARSS correction were calculated, as well as the number of greyordinates that crossed the threshold for significance as a result of MARSS correction. Spatial similarity of thresholded t-statistic maps was characterized using Sørensen–Dice coefficients^46,47^.

Finally, we quantified the change in t-statistic within an *a priori* region of interest (ROI), the left dorsolateral prefrontal cortex (lDLPFC). The lDLPFC ROI (**Figure S1**) was derived from regions of significant activation in response to SOT task demands from previous work^48,49^. The ROI was first resampled from volume to surface space, then dilated to fill holes, and finally eroded by 4mm. This analysis was performed on between-participant cortical t-statistic maps generated both with and without the use of TFCE.

## 3. Results

### 3.1 Artifact Identification and Removal Across Multiple Datasets

Elevated Pearson correlations in simultaneously acquired slices over adjacent-to-simultaneous slices across multiple task-based, *in vivo* datasets can be seen in correlation matrices in **Figure 1** (**Table S2**). Diagonal, striped patterns of elevated Pearson correlations were observed in all other *in vivo* and phantom datasets explored regardless of scanner platform or task type (**Figures S2, S3, Table S3**). Notably, the pattern of elevated correlations is directly related to the MB factor used during acquisition (see **Figure S2**). Following MARSS correction, correlations in simultaneously acquired slices were significantly reduced to values closer to the mean correlation between adjacent-to-simultaneous slices (**Table S4** for statistics).

Results of alternative artifact removal procedures tested in SBU-Siemens resting-state data are shown in **Figures S4** and **S5**. When artifact magnitude was estimated using background alone, the difference in correlation between simultaneously acquired slices and adjacent-to-simultaneous slices was reduced from 0.399 to 0.309 (W(42) = 903, p = 1.648 x 10^-8^; see **Figure S4**). This decrease in correlation, however, was much less pronounced than that produced by standard MARSS artifact correction.

Regression-based interslice leakage removal produced a slight increase in the elevation of correlations between simultaneously acquired over adjacent-to-simultaneous slices, from 0.399 to 0.405 (W(42) = 213, p = 0.029; see **Figure S5**). This indicates that no level of artifact correction was achieved by this method. **Figure S6** shows correlations between voxel pairs in simultaneously acquired slices as a function of the phase shift between the voxels during multiband reconstruction in SBU-Siemens resting-state data. Regression-based interslice leakage removal reduces the mean correlation at the expected phase shift of 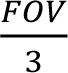 voxels (or 34 voxels) from 0.0559 in uncorrected data to -0.0273. In contrast, MARSS correction produces only a slight decrease in correlation between these voxels from 0.0559 to 0.0548.

Finally, the application of STC to the SBU-Siemens resting-state data did not produce a significant change in the difference in slice correlations between simultaneously acquired and adjacent-to-simultaneous slices (W(42) = 388, p = 0.427; see **Figure S7**).

### 3.2 Characterization of Artifact Signal

#### 3.2.1 Spatial Artifact Distribution

The spatial distributions of the average MARSS-estimated artifact signal across resting-state runs in NYSPI-GE and SBU-Siemens are shown in **Figure 2**. Images are shown on both the scale of the raw EPI data and as a percentage of the mean voxel-wise EPI signal intensity. A video depicting a 360-degree view of the rendered NYSPI-GE spatial distribution is included in the *Supplementary Information*. Artifact signal appears strongest in vasculature, notably the superior sagittal sinus, inferior sagittal sinuses, transverse sinuses, middle cerebral arteries, Circle of Willis, and superficial temporal arteries, as well as the eyes. Further, MARSS removes a larger percentage of voxel signal from both the image background and vasculature than from grey and white matter.

**Figure 2.**
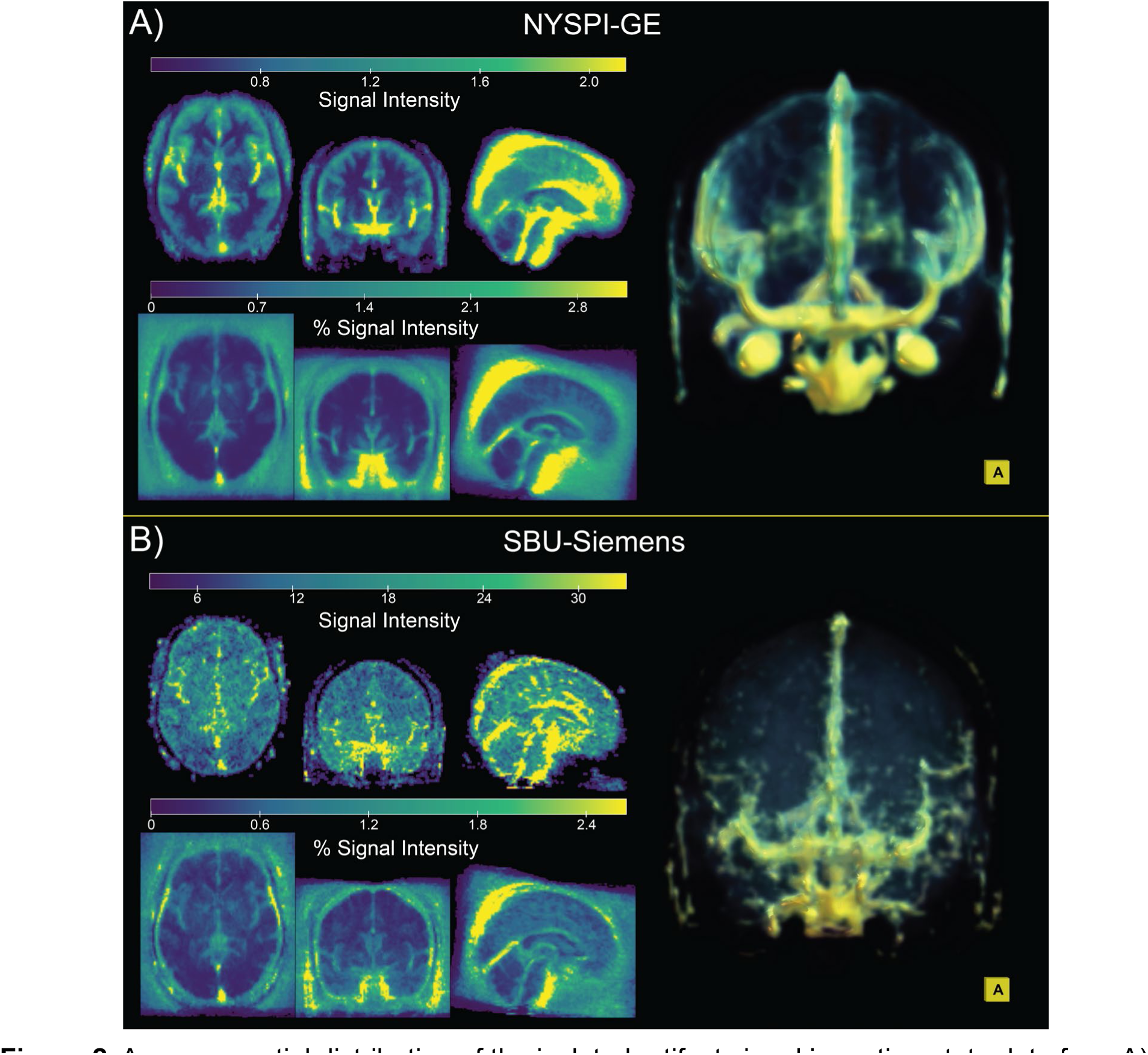
Average spatial distribution of the isolated artifact signal in resting-state data from A) NYSPI-GE (N = 63) and B) SBU-Siemens (N = 10). Artifact distribution is shown as both raw EPI signal intensity (top left of each panel) and as a percentage of the mean voxelwise EPI intensity (bottom left of each panel). Three-dimensional renders (right of each panel) display anterior view of the raw signal intensity artifact distribution.

The spatial distribution of the MARSS-estimated artifact signal in a single phantom run from SBU-Siemens acquired with a MB factor of 6 is shown in **Figure S8**. Artifact signal is strongest near the posterior edge of the phantom, where it contacted the scanner table.

#### 3.2.2 Spectral and Autocorrelative Properties

PSD estimates and autocorrelations of the MARSS-estimated artifact signal timeseries across all *in vivo*, task-based fMRI datasets are shown in **Figure 3**. All PSD estimates were linearly fit with slopes between -0.716 and -0.935 (**Figure 3**, top row). Autocorrelations in each dataset show varying degrees of long-range dependence from approximately 5 to 15 lags (**Figure 3**, bottom row). These spectral and autocorrelative characteristics roughly correspond to the properties expected of 1/f colored noise.

**Figure 3.**
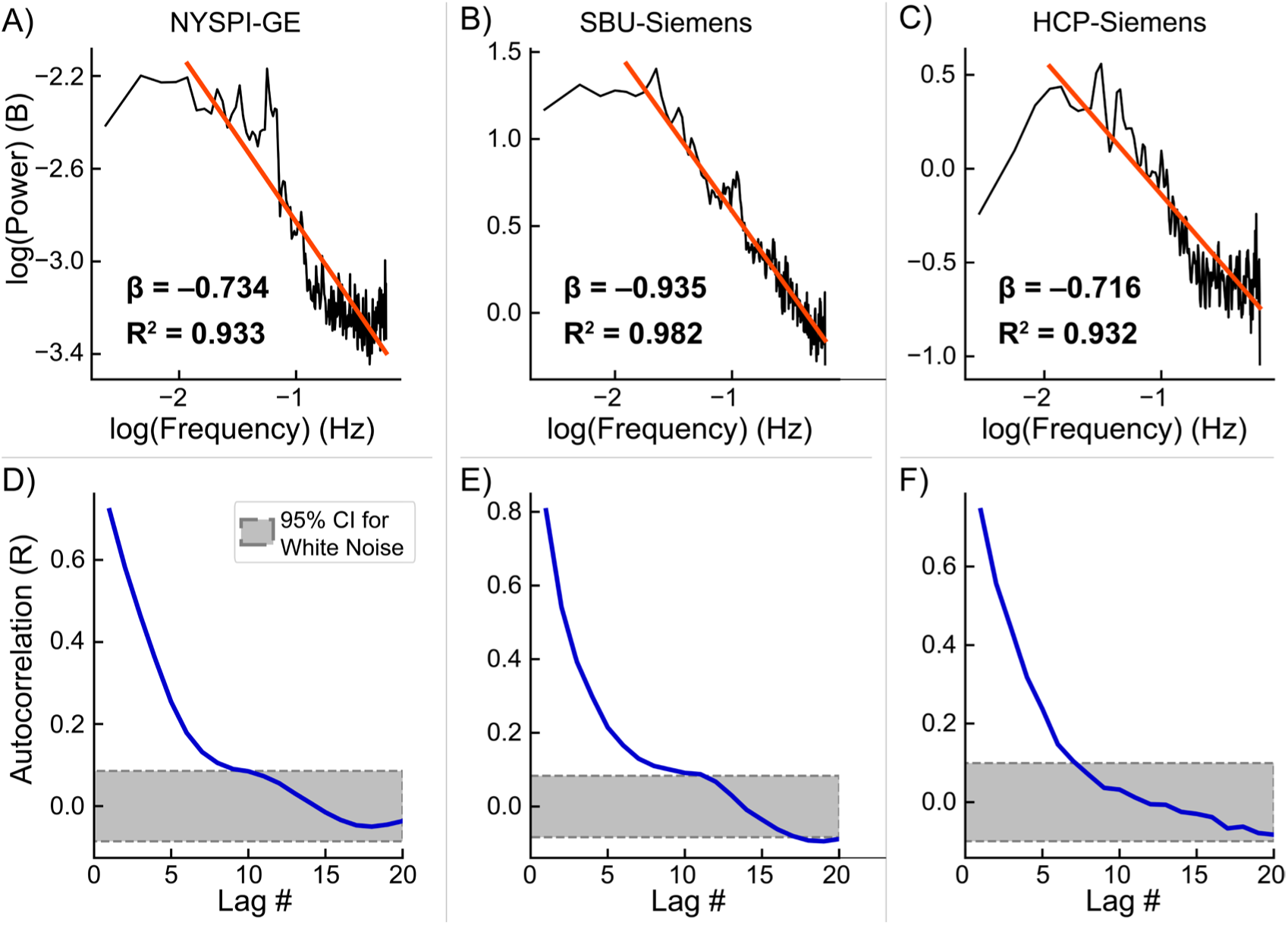
Log-log transformed power spectral density (PSD) of the isolated artifact timeseries concatenated across participants (top row) and average autocorrelation (bottom row) in task-based fMRI data from NYSPI-GE (A,D), SBU-Siemens (B,E), and HCP-Siemens (C,F). PSD plot annotations denote the slope (β) and R^2^ of each linear fit (red line). Grey areas with dotted borders in autocorrelation plots denote the 95% confidence interval for significant autocorrelation over and above that of white noise.

Additional PSD estimates are shown from all *in vivo*, resting-state fMRI datasets in **Figure S9,** and phantom data acquired with an MB factor of 6 at SBU in **Figure S10**. Autocorrelation of the artifact timeseries in *in vivo*, resting-state data as a function of lag is shown in **Figure S11.**

#### 3.2.3 Associations between Artifact Magnitude and In-Scanner Motion

Associations between mean and median measures of in-scanner motion and artifact magnitude, measured as the elevation of signal correlation between simultaneously acquired slices over adjacent-to-simultaneous slices, in NYSPI-GE resting-state and task-based data are shown in **Figure S12**. All motion measures except median DV in the NYSPI-GE resting-state dataset were significantly positively associated with artifact magnitude (all p<0.05).

#### 3.2.4 Temporal Signal-to-Noise Ratio

Across all unprocessed, *in vivo* datasets, 100% of runs showed significant increases in mean, whole-brain tSNR (all p<0.001; **Table S5**) following MARSS correction. In preprocessed, task-based data, 69.91% of voxels in NYSPI-GE and 95.16% of voxels in SBU-Siemens experienced mean tSNR increases across participants (**Figure 4**, top row; see **Figure S13** for percent change in tSNR). In preprocessed, resting-state fMRI data, 77.79% of voxels in NYSPI-GE and 97.45% of voxels in SBU-Siemens experienced mean tSNR increases (**Figure S14**; see **Figure S15** for percent change in tSNR).

**Figure 4.**
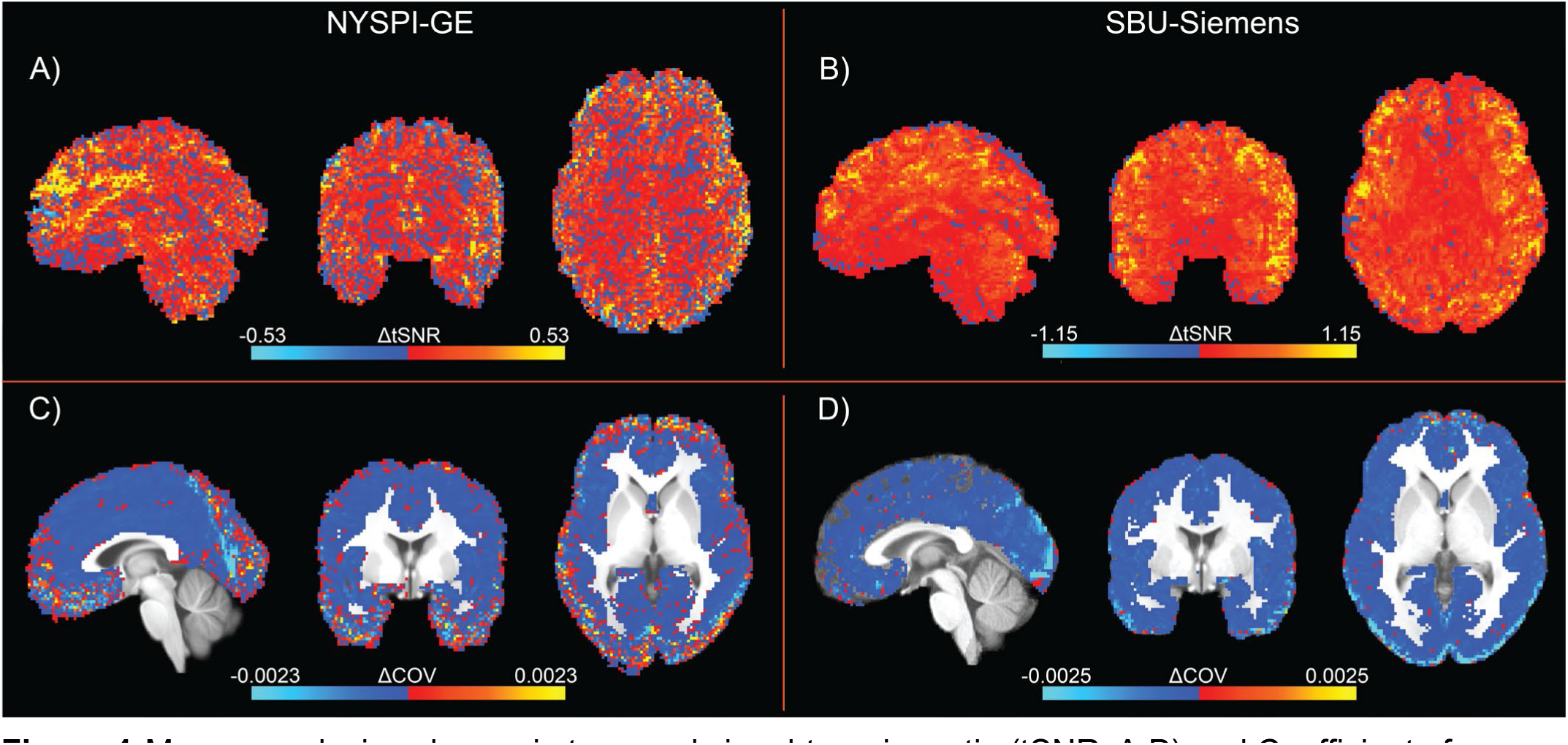
Mean, voxel-wise change in temporal signal-to-noise ratio (tSNR; A,B) and Coefficient of Variation (COV, C,D) as a result of MARSS artifact correction in preprocessed, task-based fMRI data from NYSPI-GE (A,C) and SBU-Siemens (B,D).

### 3.3 Evaluation of Artifact Magnitude in sICA+FIX Denoised Data Subspaces

The average slice correlation matrices in sICA+FIX denoised data, the neural subspace, random noise subspace, sICA+FIX noise components, and neural subspace with GSR, as well as correlation matrices for the difference between uncorrected and MARSS-corrected data, are shown in **Figure 5**. In uncorrected data, elevated correlations between simultaneously acquired slices over adjacent-to-simultaneous slices are observable in all data subspaces (all p<0.05, see **Table S6**); the correlation differences between simultaneously acquired slices and adjacent-to-simultaneous slices in sICA+FIX cleaned data and the neural subspace are 0.176 and 0.081, respectively. MARSS correction led to significant reductions in correlation differences in all subspaces (all p<0.05). Correlation matrices of the difference between uncorrected and MARSS-corrected data contain correlation elevations of simultaneously acquired slices over adjacent-to-simultaneous slices above 0.9 (see **Table S6**).

**Figure 5.**
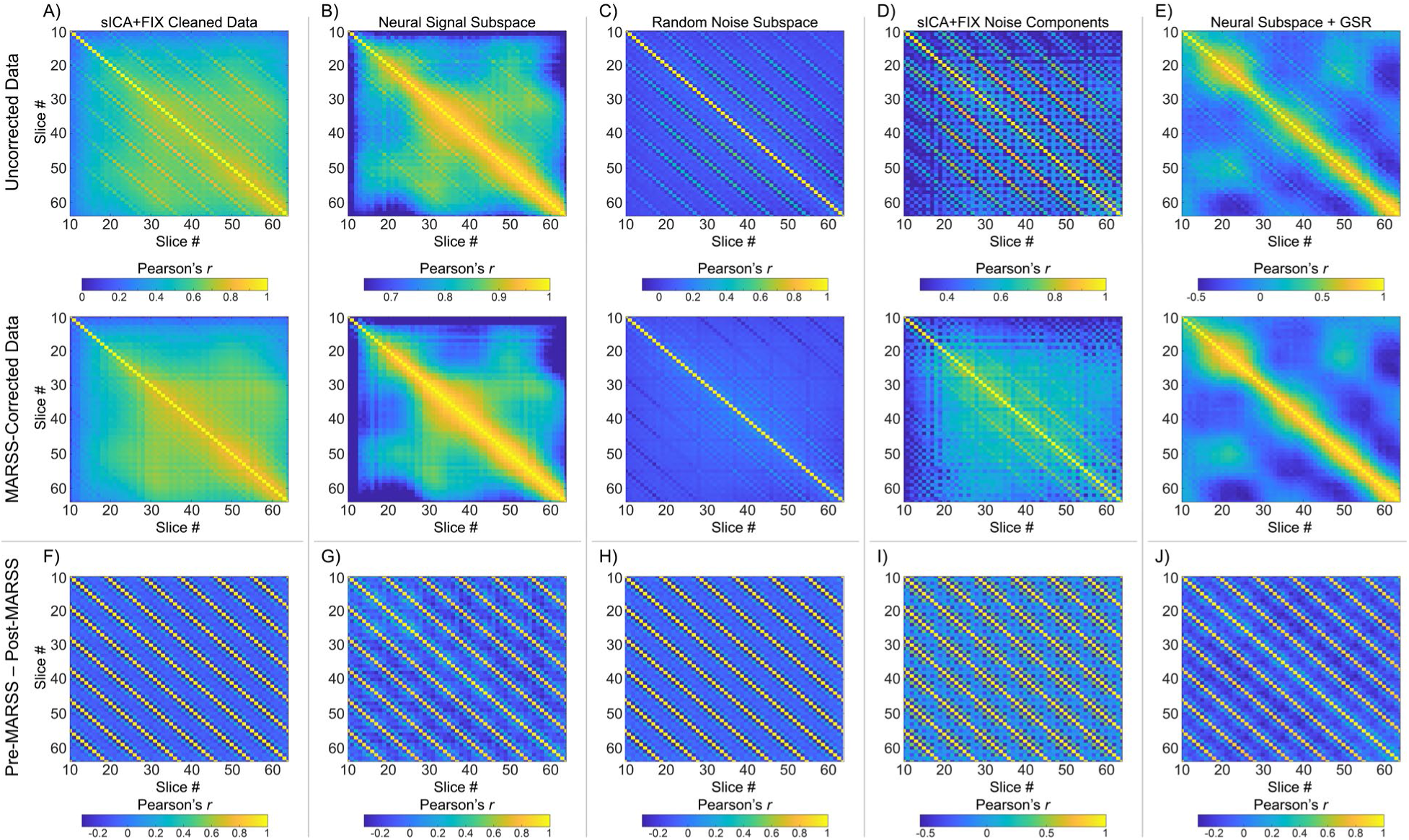
Pearson correlation matrices between average signal of all in-brain slice pairs of A) sICA+FIX cleaned data, B) the Neural Signal Subspace of sICA+FIX cleaned data, C) the Random Noise (or unstructured noise) of sICA+FIX cleaned data, D) Noise Components removed via sICA+FIX, and E) the Neural Signal Subspace after global signal regression (GSR). Correlations are only shown between slices that contain in-brain voxels across all participants. Analyses were performed in resting-state data from the HCP-Siemens dataset both prior to and following MARSS correction. Panels F-J show Pearson correlation matrices of the difference between uncorrected and MARSS-corrected data, corresponding to panels A-E, respectively.

Following MARSS correction, a statistically significant difference in correlations between simultaneously acquired slices and adjacent-to-simultaneous slices of -0.004 was observed in the neural subspace following GSR (W(100) = 1,795, p = 0.0121). Additionally, a correlation difference of 0.022 was observed in the sICA+FIX noise components (W(100) = 4,150, p = 2.307 x 10^-8^). No significant correlation differences were observed in the sICA+FIX cleaned data (W(100) = 2,165, p = 0.2158), neural signal subspace (W(100) = 1,970, p = 0.0564), and random noise subspace (W(100) = 2,382, p = 0.6229).

Results of a full variance decomposition of uncorrected and MARSS-corrected data following sICA+FIX denoising can be seen in **Table S7** and are discussed in **Supplementary Results**.

### 3.4 Within-Participants Analysis of Unprocessed Task fMRI

Single-participant examples of the difference in within-participant betas in unprocessed data from the task-on condition before and after MARSS correction are shown in **Figure 6** (top row). In all datasets, the spatial pattern of beta differences is systematic and occurs at both the fundamental spatial frequency of simultaneously acquired slice spacing and its corresponding harmonic spatial frequencies. There is a reduction in PSD at these spatial frequencies in all datasets following MARSS (**Figure 6**, bottom row).

**Figure 6.**
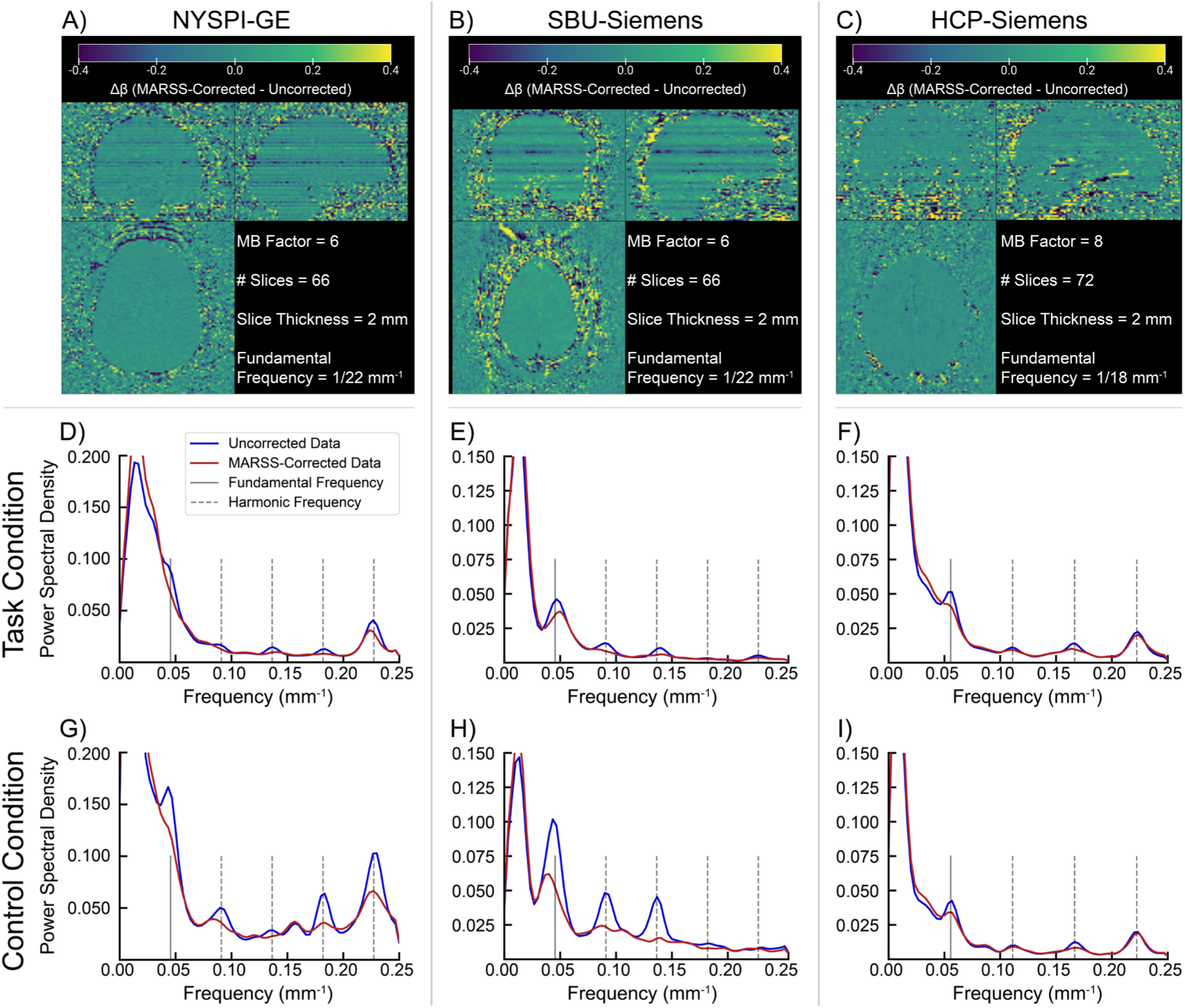
Single-participant examples of the spatial distribution of differences in task betas between MARSS-corrected and uncorrected data (A-C), as well as power spectral density (PSD) of average within-participants betas as a function of spatial frequency (D-I) in uncorrected data (blue line) and corrected data (red line) in data from NYSPI-GE (A, D, G), SBU-Siemens (B, E, H), and HCP-Siemens (F, I). In PSD plots, vertical lines denote the fundamental spatial frequency (solid grey line) and harmonic spatial frequencies (dotted grey lines) associated with the simultaneous slice pattern in each dataset. PSD is calculated separately for the task-on condition (D-F) and control condition (G-I) of each task.

### 3.5 Within- and Between-Participants Analysis of Preprocessed Task fMRI

#### 3.5.1 MARSS Correction Effects on Volume to Surface Mapping

Decreases in cortical coefficient of variation (COV) were seen in 94.62% of task-based runs in NYSPI-GE (W(186) = 534, p = 1.247 x 10^-28^) and 100% of task-based runs in SBU-Siemens (W(39) = 0, p = 5.255 x 10^-8^) following MARSS, with 80.77% of voxels in NYSPI-GE and 97.0% of voxels in SBU-Siemens experiencing mean decreases across participants (**Figure 4**, bottom row; see **Figure S13** for percent change in COV).

#### 3.5.2 MARSS Correction Effects on Within-Participants Modeling

Following MARSS, we observe a mean decrease in within-participants mean squared residuals of -0.045 (−2.16%) in NYSPI-GE (W(52) = 0, p = 3.504 x 10^-10^) and -0.062 (−5.3%) in SBU-Siemens (W(10) = 0, p = 0.002).

#### 3.5.3 Between-Participants Analysis

Figure 7 shows the difference in between-participants (second level) t-statistics obtained without TFCE before and after MARSS in NYSPI-GE and SBU-Siemens task-based data.

**Figure 7.**
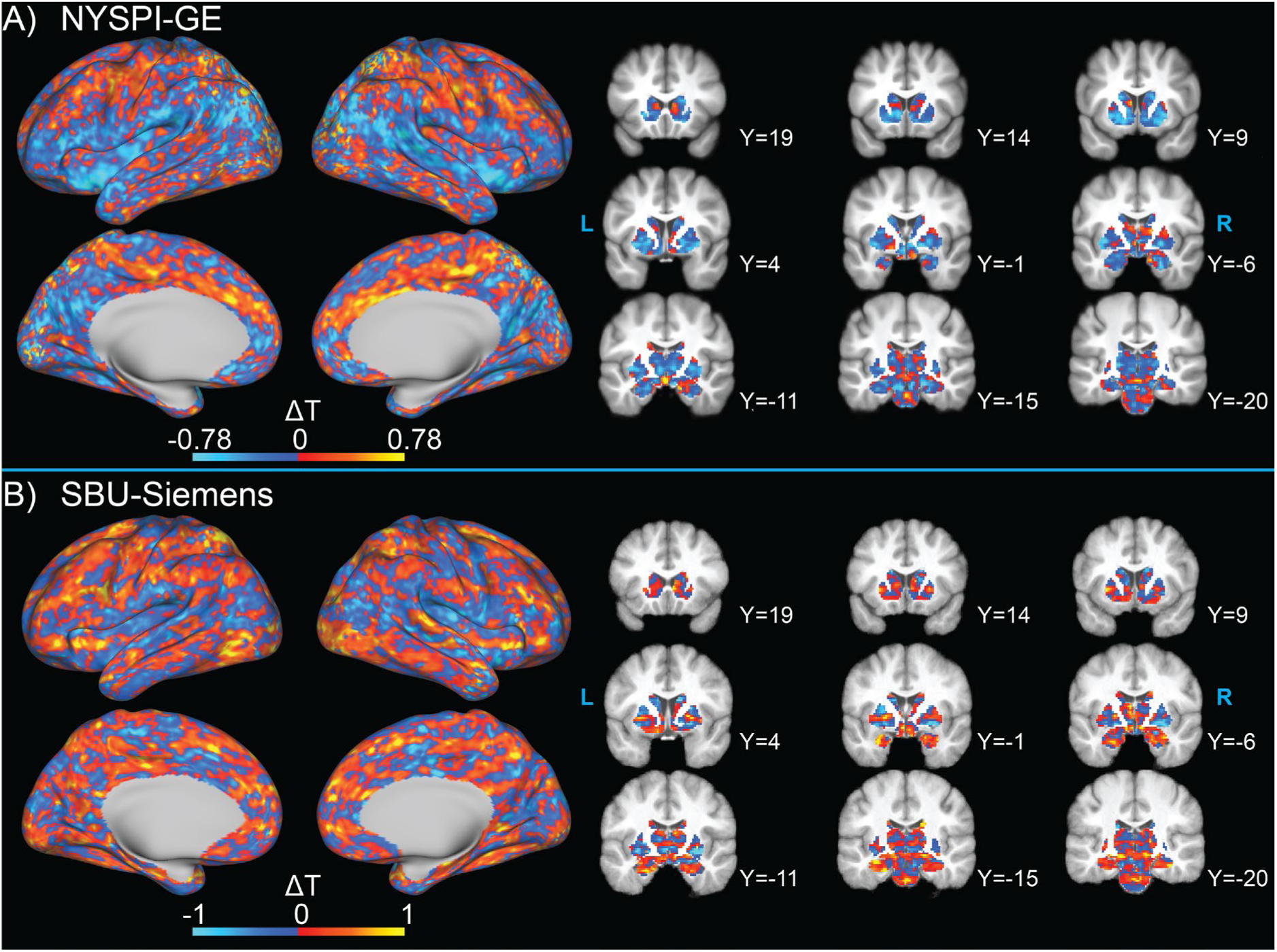
Spatial distribution of the difference in t-statistics in between-participants analyses performed in MARSS-corrected and uncorrected data from A) NYSPI-GE and B) SBU-Siemens. T-statistics were obtained without the use of threshold-free cluster enhancement (TFCE).

Striped patterns in t-statistic differences can be observed in both datasets, although these are masked by non-linear distortions in the slice pattern of the artifact, which also differ across participants, due to preprocessing. T-statistic maps reflecting the change in *magnitude* of t-statistics can be seen in **Figure S16** (if a negative t-statistic became more negative, it would be shown as a decrease in Figure 7, but as an increase in **Figure S16**). Additionally, the difference in t-statistics, as well as their magnitude difference, in between-participants analyses performed with TFCE are shown in Figure 8 and **Figure S17**, respectively. For all between-participants analyses, the deciles of absolute value of t-statistic change were calculated and are shown in **Table 2**. In NYSPI-GE, 50% of t-statistics changed by a magnitude of 0.171 or more (11.243 with TFCE), while in SBU-Siemens 50% changed by a magnitude of 0.169 or more (3.041 with TFCE).

**Figure 8.**
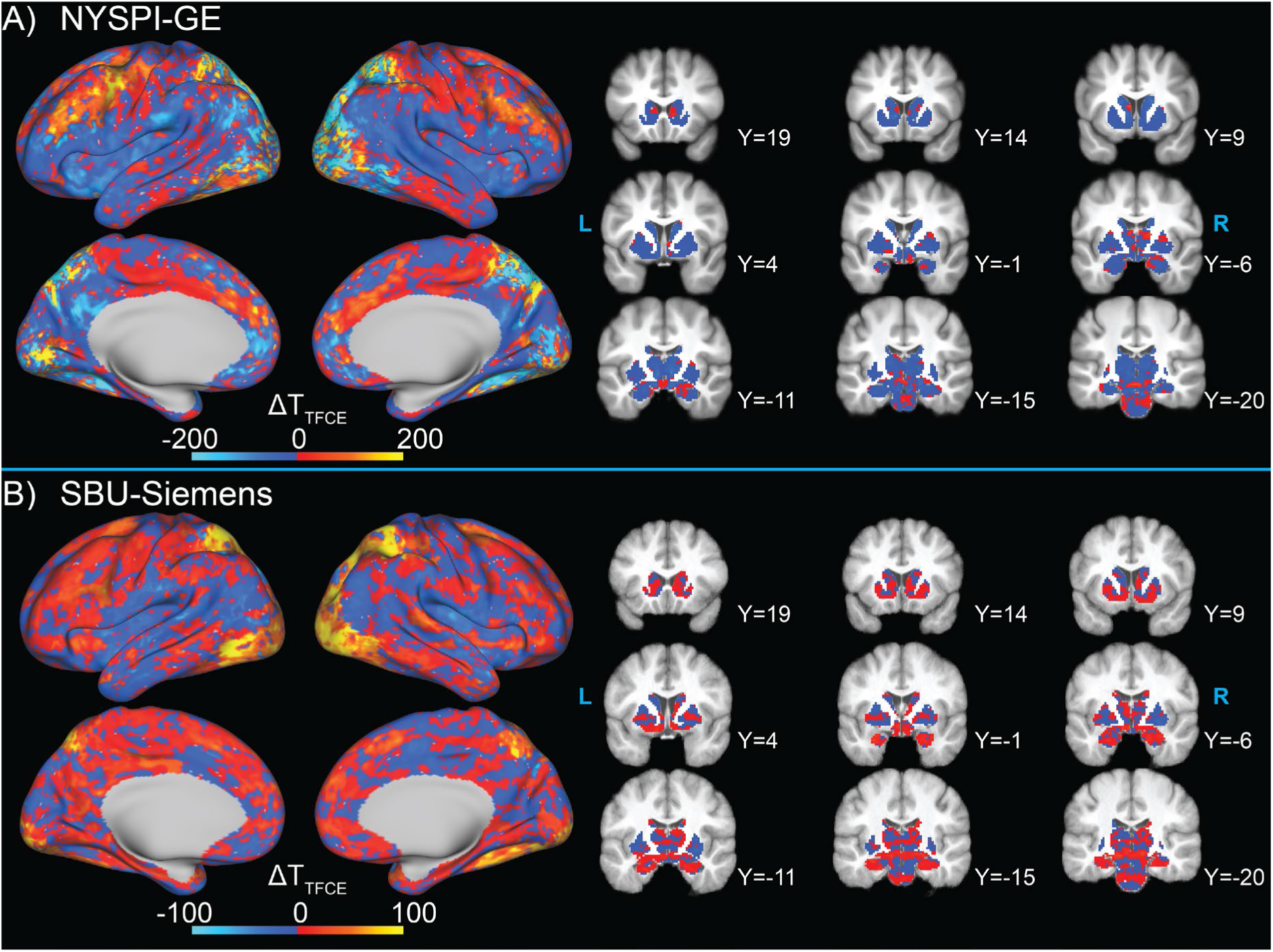
Spatial distribution of the difference in t-statistics in between-participants analysis performed in MARSS-corrected and uncorrected data from A) NYSPI-GE and B) SBU-Siemens. T-statistic were obtained using threshold-free cluster enhancement (TFCE).

**Table 2.** Deciles of the greyordinate-wise, absolute change in T-statistic resulting from MARSS artifact correction in between-participants analyses performed both with and without Threshold-free Cluster Enhancement (TFCE).

|  |  | Deciles of Absolute Change in T-statistic ("x% of T-stats change by at least _") |  |  |  |  |  |  |  |  |
| --- | --- | --- | --- | --- | --- | --- | --- | --- | --- | --- |
| Dataset | TFCE | 90% | 80% | 70% | 60% | 50% | 40% | 30% | 20% | 10% |
| NYSPI-GE | No TFCE | 0.031 | 0.063 | 0.096 | 0.132 | 0.171 | 0.216 | 0.270 | 0.341 | 0.458 |
|  | TFCE | 0.188 | 1.090 | 3.120 | 6.421 | 11.243 | 18.328 | 28.561 | 45.046 | 83.804 |
| SBU-Siemens | No TFCE | 0.030 | 0.062 | 0.094 | 0.129 | 0.169 | 0.217 | 0.275 | 0.353 | 0.490 |
|  | TFCE | 0.101 | 0.420 | 1.015 | 1.893 | 3.041 | 4.747 | 7.415 | 12.175 | 23.532 |

SBU = Stony Brook University; NYSPI = New York State Psychiatric Institute; TFCE = Threshold-free Cluster Enhancement

In NYSPI-GE, the number of significant greyordinates increased from 28,659 to 28,918 (from 40,464 to 41,458 with TFCE) as a result of MARSS correction, with 1,952 greyordinates (3,138 with TFCE) becoming significant and 1,693 (2,144 with TFCE) losing significance. In SBU-Siemens, the number of significant greyordinates increased from 8,915 to 9,828 (from 9,564 to 10,936 with TFCE), with 1,972 greyordinates (2,039 with TFCE) becoming significant and 1,059 (667 with TFCE) losing significance. Sørensen–Dice coefficients were 0.937 (0.936 with TFCE) and 0.838 (0.868 with TFCE) in NYSPI-GE and SBU-Siemens, respectively.

Within the left DLPFC ROI (**Figure S1**), the mean t-statistic increased from 5.325 to 5.376 (+0.958%) without TFCE and from 1,172.00 to 1,240.00 (+5.80%) with TFCE in NYSPI-GE. In SBU-Siemens, the mean left DLPFC t-statistic increased from 2.440 to 2.720 (+11.44%) without TFCE and from 68.231 to 83.512 (+22.40%) with TFCE.

## 4. Discussion

The results presented here demonstrate the successful mitigation of a shared artifactual signal between simultaneously acquired slices via MARSS correction. We have provided evidence that signal removed via MARSS is likely non-neural in nature, but that it impacts BOLD signal and task betas in gray matter. MARSS-corrected data exhibits higher overall tSNR and lower cortical coefficients of variation than uncorrected data. Additionally, MARSS correction mitigates residual elevations in correlations between simultaneously acquired slices after denoising via sICA+FIX. In task-based analyses, MARSS results in decreased participant-level task model error and removes spatial patterns of task betas at spatial frequencies corresponding to the distance between simultaneously acquired slices. Between participants, MARSS causes substantive changes in group-level t-statistics that tend to follow striped patterns along the slice acquisition plane, even after spatial normalization is performed. These results collectively indicate significant improvements in data quality following MARSS correction.

### 4.1 MARSS Correction Reduces Simultaneous Slice Correlations

We have shown the presence of a shared artifactual signal between simultaneously acquired slices of multiband fMRI data acquired across six datasets, including in two widely used publicly available datasets^7,9^. This signal was originally identified as elevated correlations between timeseries in simultaneously acquired slices relative to adjacent-to-simultaneous slices, in both phantom and *in vivo* data, with elevations exceeding a difference in Pearson *r* correlation coefficients of 0.3 or 0.4 in many datasets. Further, the spatial pattern of elevated correlations follows a clear, systematic relationship with the MB factor used during acquisition. As there is no physiological justification for BOLD signal in arbitrary axial slices to be more related in simultaneously acquired slices than in any other given pair of slices (including neighboring slices), we developed a regression-based detection and correction technique, MARSS, that estimates the shared signal across simultaneously acquired slices in unprocessed MB-fMRI data and removes the estimated signal from each voxel in an fMRI run. This technique successfully reduces the magnitude of the difference in correlations between simultaneously acquired and adjacent-to-simultaneous slices.

Although MARSS also reduced simultaneous slice correlations to levels slightly below that of the average correlation between adjacent-to-simultaneous slices, it does so to a much lesser degree than the magnitude of the original elevated correlations. As shown in **Table S2**, the maximum negative difference in correlation between simultaneously acquired slices and adjacent-to-simultaneous slices induced by MARSS correction in any *in vivo* dataset is -0.085, as opposed to 0.465 in uncorrected data. Although we collected only minimal data at lower MB factors (e.g., 3 or 4), the data in **Table S3** raise the possibility that lower MB factors are associated with smaller magnitude reductions in simultaneous-slice correlations after MARSS correction. This is consistent with the fact that lower MB factors should, *a priori*, produce poorer estimates of the shared signal across slices due to having fewer slices to average together to arrive at an estimate of the artifact.

### 4.2 MARSS Correction Improves SNR and Removes Non-Neural Signal

Critically, we show that MARSS artifact correction results in the removal of image noise rather than true neural signal. First, an increase in mean, whole-brain tSNR in 100% of unprocessed runs across all datasets after MARSS correction suggests improved data quality. In preprocessed data, the largest mean increases in tSNR occurred in cortical regions, while many decreases occurred at tissue boundaries where BOLD signal most likely contained greater contamination from fluid motion or vascular signal (Figure 4, top row; **Figures S13-15**). Although tSNR carries no information on activation strength in task-based fMRI data, it is a useful measure of signal stability in studies of resting-state fMRI^50^.

One major indication that MARSS predominantly removes non-neural signal lies in the spatial distribution of the estimated artifact in Figure 2, where the largest amount of signal is removed from large vasculature, and a higher percentage of mean signal is removed from the background of the image. The significant vascular involvement suggests that artifact correction is isolating signal related to vasomotion or pulsatile blood flow whose contribution elevates signal correlations between simultaneously acquired slices. It is plausible that this signal originates from systemic low-frequency oscillations (sLFOs), which are considered non-neural in nature and account for a significant portion of the variance in signal in the range of approximately 0.01-0.15 Hz^51,52^. Notably, isolated artifact timeseries show autocorrelations up to lags that approximately correspond with this frequency range (Figure 3 and **Figure S11**).

The frequency band occupied by sLFOs is the same as that which is occupied by fluctuations in neural activity that are the basis of resting-state functional connectivity. Thus, traditional spectral filtering techniques cannot remove these signals without compromising true signal from neuronal fluctuations. Tong and colleagues demonstrated that sLFOs can confound estimates of resting-state functional connectivity, going as far as mimicking resting-state networks isolated via independent components analysis (ICA)^53^. Additionally, they have identified major vascular fMRI signal components that appear in the spatial distribution of the artifact signal, such as the superior sagittal sinus, to be highly correlated with simultaneously-acquired near infrared spectroscopy estimates of blood flow^54^.

Slightly complicating this explanation of the artifact source is its presence in phantom data. The artifact signal intensity in phantom data is strongest near the point of contact with the scanner table (see **Figure S8**). Due to this being a liquid-filled phantom, it is possible that scanner table vibrations caused periodic fluid motion that resembles pulsatile blood flow. While currently inconclusive, the possibility that MARSS artifact estimation is partially or completely identifying signal originating from the vasculature bears further investigation.

Given that MARSS estimates and corrects for a structured noise signal that relies on the simultaneous slice acquisition pattern in unprocessed scanner space for identification, denoising techniques such as sICA+FIX that are typically performed on data that has been normalized to a template space, which distorts the slice acquisition pattern, would not be expected to fully account for this signal. This is evident in the correlation matrices of resting-state data from the HCP-Siemens dataset that has been denoised using sICA+FIX and decomposed into neural and random noise subspaces (see Figure 5). Prior to MARSS correction, elevated correlations between simultaneously acquired slices show up strongly in both subspaces, indicating that a portion of the artifact signal was likely modeled as true neural signal across multiple ICs, and the remaining artifact signal in the cleaned data was not modeled. Further, GSR is not able to correct the elevated correlations in the neural subspace, instead slightly exacerbating them. In contrast, MARSS correction prior to sICA+FIX denoising significantly reduces artifact presence across all subspaces, resulting in statistically insignificant correlation differences between simultaneously acquired slices and adjacent-to-simultaneous slices. Correlation matrices of the difference between uncorrected and MARSS-corrected data do not exhibit any structured correlation patterns besides that between simultaneously acquired slices, indicating that MARSS correction is most likely not removing neural signal from any subspace. While further evaluation of sICA+FIX denoising in MARSS-corrected fMRI data is needed, these results suggest that MARSS and sICA+FIX denoising used in tandem may provide maximal denoising of structured artifact signals.

### 4.3 MARSS Correction Reduces Cortical Coefficient of Variation

The removal of noise from unprocessed, volumetric data due to artifact correction is beneficial to volume-to-surface mapping in the HCP MPP^36^. For context, the mapping algorithm excludes voxels with coefficients of variation that are 0.5 standard deviations above the mean local coefficient of variation in a 5mm Gaussian neighborhood. Both the NYSPI-GE and SBU-Siemens datasets experience decreases in coefficient of variation in 97.00% and 80.77% of voxels, respectively. This nearly global decrease suggests an increase in probability of most cortical voxels to be included in volume to surface mapping. Notably, several runs of NYSPI-GE data were discovered to have striped patterns of high coefficients of variation (see **Figure S22**) that are mitigated after MARSS correction. Upon inspection, data from these runs show no discernable quality control concerns, suggesting that multiband acquisition can sometimes cause spurious, yet systematic noise patterns that could potentially surface timeseries extraction in otherwise usable data.

### 4.4 MARSS Correction Improves Spatial Statistics Within and Between Participants

Critically, we have shown that MARSS correction has substantial implications for spatial statistics in task-based analyses, both within and between participants. When comparing within-participants analyses in unprocessed data prior to and following MARSS correction, we observe a striping pattern in task beta differences across all task-based datasets (Figure 6, top row). We analyzed the spatial frequency of these patterns along the slice direction and demonstrated that systematic patterns in task betas occur at the fundamental frequency and harmonic frequencies of the simultaneously acquired slice pattern in uncorrected data (Figure 6, bottom two rows), and that the spectral power at these spatial frequencies is reduced by MARSS. While there is no ground truth expectation as to the magnitude of task betas, it is reasonable to conclude that true estimates of task-evoked neuronal activation should not exhibit a spatial frequency dependent on the spacing between simultaneously acquired slices. The fact that MARSS correction reduces the spectral power of these patterns indicates that this spatial frequency is being removed from the data by our method, rather than being added.

In within-participants analyses of preprocessed, task-based data, we observe statistically significant decreases in mean-squared residuals, indicating lower model error and suggesting that task-irrelevant, and potentially non-neural, signals are being removed by MARSS. At the group level, we observe striping patterns in the difference in t-statistics between analyses performed before and after MARSS correction (Figures 7**, 8**). Spatial frequency analyses demonstrated that the systematic pattern is present first in unprocessed, uncorrected data and persists in the data even after spatial normalization and volume-to-surface resampling distort the slice acquisition pattern to varying degrees in each individual participant. MARSS correction led to substantial changes in between-participants t-statistics, with 10% of greyordinates changing by at least 0.458 and 0.490 in NYSPI-GE and SBU-Siemens, respectively (**Table 2**).

This resulted in a net increase in the number of significant greyordinates, both with and without the use of TFCE. Sørensen–Dice coefficients for each dataset indicate a change in significance in approximately 6.63% (6.64% with TFCE) greyordinates in NYSPI-GE and 16.20% (13.20% with TFCE) greyordinates in SBU-Siemens. While high Sørensen–Dice coefficients are typically desirable in the validation of image segmentation algorithms and other deep learning applications, a lower coefficient in this application indicates a greater change in spatial statistics induced by MARSS correction.

Since there is no ground truth expectation for estimations of whole-brain, task-evoked activation, we then constrained our analysis to within the left DLPFC, which is known to show robust activation in response to working memory task demands^55,56,57^. Within the chosen ROI, which was derived from previous group-level analysis of the SOT in a separate dataset^48,49^, t-statistics increase in both NYSPI-GE and SBU-Siemens with and without the use of TFCE. The larger percentage increase in t-statistic that results from using TFCE suggests that removal of noise due to artifact correction enhances the spatial homogeneity of patterns of true task-evoked activation, leading to clearer identification and amplification of significant regions by the TFCE algorithm.

### 4.5 Differences in Results Between Datasets

One theme in the results presented here is that NYSPI-GE appears to show less improvement than SBU-Siemens in between-participants analyses following MARSS correction. There are several hardware differences between these datasets that may influence this, such as scanner manufacturer and coil configuration. Further, the slightly longer TR used in NYSPI-GE data could result in diminished temporal resolution and less adequate sampling of artifactual periodic signals identified by artifact estimation. Despite these differences, the spatial distribution of isolated artifact signal (Figure 2) in NYSPI-GE is highly similar to that of SBU-Siemens; differences in smoothness of the spatial distribution can be attributed to the smaller sample size of SBU-Siemens (N =10) relative to that of NYSPI-GE (N = 63). Additionally, both datasets experience improvements in tSNR and reductions in cortical COV, as well as reduction of artificially elevated PSD of task betas at fundamental and harmonic frequencies related to the simultaneous slice acquisition, indicating improvement in task modeling regardless of the resulting change in between-participants t-statistics.

### 4.6 Limitations and Future Considerations

This study establishes the foundation for several analyses to further investigate the source of the observed artifact signal that have yet to be conducted. First, the true impact of scanner manufacturer or platform would best be investigated using an identical phantom, and potentially human participants, and acquisition sequence across multiple scanners. Within this analysis, scanner vibration^58^ should be recorded to assess the impact of mechanical noise on artifact magnitude. Next, acquisition parameters such as partial Fourier sampling factor and phase-encode direction should be varied in order to determine if there are optimal combinations of parameters that minimize correlation elevations between simultaneously acquired slices.

Additionally, the effect of variations in raw data reconstruction pipeline should be assessed, including an evaluation of artifact presence in data reconstructed using slice-GRAPPA^5^ and split-slice GRAPPA^11^.

While we have demonstrated that a shared artifact signal between simultaneously acquired slices is present even after denoising via sICA+FIX, subsequent analyses should include an evaluation of MARSS correction used both in tandem with and against other existing denoising techniques, such as temporal ICA^15,59^, ICA AROMA^60^, RETROICOR^61^, and ANATICOR^62,63^. The collection of cardiac and respiratory traces is necessary for the implementation of RETROICOR, and spectral characteristics and cross-correlations between these physiological signals and the artifact isolated by MARSS correction should be investigated.

Furthermore, the sICA+FIX denoising analyses presented here used ICs estimated on uncorrected data, but then applied to MARSS-corrected data. A full examination of how MARSS and sICA+FIX or other ICA-based methods interact would require the estimation of ICA components on MARSS-corrected data. These analyses should include a comparison of both estimates of task-evoked activation and resting-state functional connectivity across techniques.

An additional limitation is that our examination of MARSS denoising here is primarily restricted to MB acceleration factors of 6 and 8. As noted in Section 4.1 above, lower MB acceleration factors should be expected to produce poorer estimates of the artifact signal (due to the existence of fewer simultaneously-acquired slices to estimate the artifact from), and there is no guarantee that our results would hold at lower MB factors. At a minimum, application of MARSS should probably not be conducted at an MB factor of 2, as this would effectively involve estimating the MARSS artifact from only a single slice, with no averaging of spatially disparate slices that helps to mitigate the impact of true underlying neural signals on the estimated artifact signal.

### 4.7 Conclusion

In summary, we have demonstrated the presence and successful correction of an artifactual, non-neural signal that is shared between simultaneously acquired slices in multiple datasets. Removal of this artifact with the MARSS method developed here results in substantive improvements in data quality, including nearly global increases in tSNR and improvements to the extraction of the cortical surface during preprocessing. The artifact appears strongest in the neurovasculature, but has clearly demonstrable impacts throughout grey matter voxels and greyordinates. Further, the artifact is not fully accounted for by sICA+FIX denoising, likely because ICA components are estimated after spatial distortions of the raw scanner space data have occurred during preprocessing, thereby mixing artifact signals from adjacent-to-simultaneous slices together and making the true original MARSS artifact signal unrecoverable following standard preprocessing. In addition, MARSS correction leads to a reduction of systematic spatial patterns in estimates of task-evoked activation related to slice acquisition patterns, both within and between participants, which ultimately leads to substantive changes in group-level t-statistics throughout the brain. We expect that application of our artifact correction method will lead to an improvement in both the quality and reproducibility of all multiband fMRI datasets, and recommend that investigators implement this technique prior to functional data preprocessing, at least for MB factors of 6 or higher.

## 5. Acknowledgements and Disclosures

### 5.2 Acknowledgements

The authors would like to thank Jack Grinband for comments and suggestions on preliminary findings in this work, and Matthew F. Glasser for guidance and feedback on the implementation and evaluation of sICA+FIX denoising. We thank Yash Patel, Jaeyop Jeong, Sam R. Luceno, and Sameera Abeykoon for their contributions to fMRI data preprocessing and Mahika Gupta for assistance with preparing MARSS code for public release. Next, we would like to thank Guarav Patel and Matthew Riddle at the New York State Psychiatric Institute, and Turhan Canli and Kim Burke at Stony Brook University for their assistance in acquiring phantom data. Finally we would like to acknowledge the substantial computing resources and technical assistance provided by Stony Brook Medicine Research Computing, with substantial support from Allen Zawada and James Xikis, as well as Stony Brook Research Computing and Cyberinfrastructure and the Institute for Advanced Computational Science at Stony Brook University for access to the high-performance SeaWulf computing system (National Science Foundation Award Nos. 1531492 and 2215987, and matching funds from the Empire State Development’s Division of Science, Technology and Innovation program contract C210148), with notable support from Fırat Coşkun, Daniel Wood, and David Carlson.

## 6. Data Availability

Data used in this manuscript can be made available by the authors upon request.

Publicly available HCP data can be obtained from ConnectomeDB (https://db.humanconnectome.org/)^39^. ABCD study data can be obtained from the National Institute of Mental Health National Data Archive (NIMH NDA).

## 7. Code Availability

MARSS is available as an open-source MATLAB software package on GitHub (https://github.com/CNaP-Lab/MARSS) and the MathWorks File Exchange (https://www.mathworks.com/matlabcentral/fileexchange/156782-multiband-artifact-regression-in-simultaneous-slices-marss). Code to replicate all analyses in this manuscript can be made available by the authors upon request.

## References

1. Feinberg DA, et al. Multiplexed echo planar imaging for sub-second whole brain FMRI and fast diffusion imaging. PLoS One 5, e15710 (2010).

2. Moeller S, et al. Multiband multislice GE-EPI at 7 tesla, with 16-fold acceleration using partial parallel imaging with application to high spatial and temporal whole-brain fMRI. Magn Reson Med 63, 1144–1153 (2010).

3. Xu J, et al. Evaluation of slice accelerations using multiband echo planar imaging at 3 T. Neuroimage 83, 991–1001 (2013).

4. Breuer FA, Blaimer M, Heidemann RM, Mueller MF, Griswold MA, Jakob PM. Controlled aliasing in parallel imaging results in higher acceleration (CAIPIRINHA) for multi-slice imaging. Magnetic Resonance in Medicine 53, 684–691 (2005).

5. Setsompop K, Gagoski BA, Polimeni JR, Witzel T, Wedeen VJ, Wald LL. Blipped-controlled aliasing in parallel imaging for simultaneous multislice echo planar imaging with reduced g-factor penalty. Magnetic Resonance in Medicine 67, 1210–1224 (2011).

6. Ugurbil K, et al. Pushing spatial and temporal resolution for functional and diffusion MRI in the Human Connectome Project. Neuroimage 80, 80–104 (2013).

7. Van Essen DC, et al. The WU-Minn Human Connectome Project: an overview. Neuroimage 80, 62–79 (2013).

8. Miller KL, et al. Multimodal population brain imaging in the UK Biobank prospective epidemiological study. Nat Neurosci 19, 1523–1536 (2016).

9. Casey BJ, et al. The Adolescent Brain Cognitive Development (ABCD) study: Imaging acquisition across 21 sites. Dev Cogn Neurosci 32, 43–54 (2018).

10. Todd N, Moeller S, Auerbach EJ, Yacoub E, Flandin G, Weiskopf N. Evaluation of 2D multiband EPI imaging for high-resolution, whole-brain, task-based fMRI studies at 3T: Sensitivity and slice leakage artifacts. Neuroimage 124, 32–42 (2016).

11. Cauley SF, Polimeni JR, Bhat H, Wald LL, Setsompop K. Interslice leakage artifact reduction technique for simultaneous multislice acquisitions. Magnetic Resonance in Medicine 72, 93–102 (2014).

12. Griswold MA, et al. Generalized autocalibrating partially parallel acquisitions (GRAPPA). Magnetic Resonance in Medicine 47, 1202–1210 (2002).

13. McNabb CB, Lindner M, Shen S, Burgess LG, Murayama K, Johnstone T. Inter-slice leakage and intra-slice aliasing in simultaneous multi-slice echo-planar images. Brain Struct Funct 225, 1153–1158 (2020).

14. Burgess GC, et al. Evaluation of Denoising Strategies to Address Motion-Correlated Artifacts in Resting-State Functional Magnetic Resonance Imaging Data from the Human Connectome Project. Brain Connectivity 6, 669–680 (2016).

15. Glasser MF, et al. Using temporal ICA to selectively remove global noise while preserving global signal in functional MRI data. NeuroImage 181, 692–717 (2018).

16. Power JD, Lynch CJ, Silver BM, Dubin MJ, Martin A, Jones RM. Distinctions among real and apparent respiratory motions in human fMRI data. NeuroImage 201, (2019).

17. Fair DA, et al. Correction of respiratory artifacts in MRI head motion estimates. NeuroImage 208, (2020).

18. Williams JC, Tubiolo PN, Luceno JR, Van Snellenberg JX. Advancing motion denoising of multiband resting-state functional connectivity fMRI data. Neuroimage 249, 118907 (2022).

19. Power JD, Lynch CJ, Silver BM, Dubin MJ, Martin A, Jones RM. Distinctions among real and apparent respiratory motions in human fMRI data. Neuroimage 201, 116041 (2019).

20. Fair DA, et al. Correction of respiratory artifacts in MRI head motion estimates. Neuroimage 208, 116400 (2020).

21. Burgess GC, et al. Evaluation of Denoising Strategies to Address Motion-Correlated Artifacts in Resting-State Functional Magnetic Resonance Imaging Data from the Human Connectome Project. Brain Connect 6, 669–680 (2016).

22. Power JD, Mitra A, Laumann TO, Snyder AZ, Schlaggar BL, Petersen SE. Methods to detect, characterize, and remove motion artifact in resting state fMRI. NeuroImage 84, 320–341 (2014).

23. Griffanti L, et al. ICA-based artefact removal and accelerated fMRI acquisition for improved resting state network imaging. NeuroImage 95, 232–247 (2014).

24. Salimi-Khorshidi G, Douaud G, Beckmann CF, Glasser MF, Griffanti L, Smith SM. Automatic denoising of functional MRI data: Combining independent component analysis and hierarchical fusion of classifiers. NeuroImage 90, 449–468 (2014).

25. Van Snellenberg JX, et al. Dynamic shifts in brain network activation during supracapacity working memory task performance. Human Brain Mapping 36, 1245–1264 (2015).

26. Barch DM, et al. Function in the human connectome: Task-fMRI and individual differences in behavior. NeuroImage 80, 169–189 (2013).

27. Friedman L, Glover GH. Report on a multicenter fMRI quality assurance protocol. Journal of Magnetic Resonance Imaging 23, 827–839 (2006).

28. Gibbons JD, Chakraborti S. Nonparametric statistical inference, 6th edition. edn. CRC Press (2021).

29. Hollander M, Wolfe DA, Chicken E. Nonparametric statistical methods, Third edition / edn. John Wiley & Sons, Inc. (2014).

30. Hoopes A, Mora JS, Dalca AV, Fischl B, Hoffmann M. SynthStrip: skull-stripping for any brain image. NeuroImage 260, (2022).

31. Fischl B. FreeSurfer. Neuroimage 62, 774–781 (2012).

32. Welch P. The use of fast Fourier transform for the estimation of power spectra: A method based on time averaging over short, modified periodograms. IEEE Transactions on Audio and Electroacoustics 15, 70–73 (1967).

33. Friston KJ. Statistical parametric mapping: the analysis of funtional brain images, 1st edn. Elsevier/Academic Press (2007).

34. Friston KJ, Holmes AP, Worsley KJ, Poline JP, Frith CD, Frackowiak RSJ. Statistical parametric maps in functional imaging: A general linear approach. Human Brain Mapping 2, 189–210 (2004).

35. Grabner G, Janke AL, Budge MM, Smith D, Pruessner J, Collins DL. Symmetric Atlasing and Model Based Segmentation: An Application to the Hippocampus in Older Adults. In: Medical Image Computing and Computer-Assisted Intervention – MICCAI 2006) (2006).

36. Glasser MF, et al. The minimal preprocessing pipelines for the Human Connectome Project. NeuroImage 80, 105–124 (2013).

37. Murphy K, Bodurka J, Bandettini PA. How long to scan? The relationship between fMRI temporal signal to noise ratio and necessary scan duration. NeuroImage 34, 565–574 (2007).

38. Welvaert M, Rosseel Y. On the Definition of Signal-To-Noise Ratio and Contrast-To-Noise Ratio for fMRI Data. PLoS ONE 8, (2013).

39. Hodge MR, et al. ConnectomeDB—Sharing human brain connectivity data. NeuroImage 124, 1102–1107 (2016).

40. Winkler AM, Ridgway GR, Webster MA, Smith SM, Nichols TE. Permutation inference for the general linear model. NeuroImage 92, 381–397 (2014).

41. Winkler AM, Ridgway GR, Douaud G, Nichols TE, Smith SM. Faster permutation inference in brain imaging. NeuroImage 141, 502–516 (2016).

42. Winkler AM, Webster MA, Brooks JC, Tracey I, Smith SM, Nichols TE. Non-parametric combination and related permutation tests for neuroimaging. Human Brain Mapping 37, 1486–1511 (2016).

43. Winkler AM, Webster MA, Vidaurre D, Nichols TE, Smith SM. Multi-level block permutation. Neuroimage 123, 253–268 (2015).

44. Smith S, Nichols T. Threshold-free cluster enhancement: Addressing problems of smoothing, threshold dependence and localisation in cluster inference. NeuroImage 44, 83–98 (2009).

45. Šidák Z. Rectangular Confidence Regions for the Means of Multivariate Normal Distributions. Journal of the American Statistical Association 62, 626–633 (1967).

46. Dice LR. Measures of the Amount of Ecologic Association Between Species. Ecology 26, 297–302 (1945).

47. Sørensen TJ. A method of establishing groups of equal amplitude in plant sociology based on similarity of species content and its application to analyses of the vegetation on Danish commons. København: I kommission hos E. Munksgaard, 1948. (1948).

48. Van Snellenberg JX, et al. Dynamic shifts in brain network activation during supracapacity working memory task performance. Hum Brain Mapp 36, 1245–1264 (2015).

49. Van Snellenberg JX, et al. Mechanisms of Working Memory Impairment in Schizophrenia. Biol Psychiatry 80, 617–626 (2016).

50. De Blasi B, et al. Noise removal in resting-state and task fMRI: functional connectivity and activation maps. Journal of Neural Engineering 17, (2020).

51. Frederick Bd, Nickerson LD, Tong Y. Physiological denoising of BOLD fMRI data using Regressor Interpolation at Progressive Time Delays (RIPTiDe) processing of concurrent fMRI and near-infrared spectroscopy (NIRS). NeuroImage 60, 1913–1923 (2012).

52. Tong Y, Hocke LM, Frederick BB. Low Frequency Systemic Hemodynamic “Noise” in Resting State BOLD fMRI: Characteristics, Causes, Implications, Mitigation Strategies, and Applications. Frontiers in Neuroscience 13, (2019).

53. Tong Y, Hocke LM, Fan X, Janes AC, Frederick Bd. Can apparent resting state connectivity arise from systemic fluctuations? Frontiers in Human Neuroscience 9, (2015).

54. Tong Y, Hocke LM, Nickerson LD, Licata SC, Lindsey KP, Frederick Bd. Evaluating the effects of systemic low frequency oscillations measured in the periphery on the independent component analysis results of resting state networks. NeuroImage 76, 202–215 (2013).

55. Barbey AK, Koenigs M, Grafman J. Dorsolateral prefrontal contributions to human working memory. Cortex 49, 1195–1205 (2013).

56. Funahashi S, Takeda K, Watanabe Y. Neural mechanisms of spatial working memory: Contributions of the dorsolateral prefrontal cortex and the thalamic mediodorsal nucleus. *Cognitive, Affective*, & Behavioral Neuroscience 4, 409–420 (2004).

57. Hampson M, Zhang G, Yao L, Zhang H, Long Z, Zhao X. Improved Working Memory Performance through Self-Regulation of Dorsal Lateral Prefrontal Cortex Activation Using Real-Time fMRI. PLoS ONE 8, (2013).

58. Berl MM, et al. Investigation of vibration-induced artifact in clinical diffusion-weighted imaging of pediatric subjects. Human Brain Mapping 36, 4745–4757 (2015).

59. Glasser MF, et al. Classification of temporal ICA components for separating global noise from fMRI data: Reply to Power. NeuroImage 197, 435–438 (2019).

60. Pruim RHR, Mennes M, van Rooij D, Llera A, Buitelaar JK, Beckmann CF. ICA-AROMA: A robust ICA-based strategy for removing motion artifacts from fMRI data. NeuroImage 112, 267–277 (2015).

61. Glover GH, Li T-Q, Ress D. Image-based method for retrospective correction of physiological motion effects in fMRI: RETROICOR. Magnetic Resonance in Medicine 44, 162–167 (2000).

62. Jo HJ, et al. Fast detection and reduction of local transient artifacts in resting-state fMRI. Computers in Biology and Medicine 120, (2020).

63. Jo HJ, Saad ZS, Simmons WK, Milbury LA, Cox RW. Mapping sources of correlation in resting state FMRI, with artifact detection and removal. NeuroImage 52, 571–582 (2010).

