## Supplementary Material for "Characterization and Mitigation of a Simultaneous Multi-Slice fMRI Artifact: Multiband Artifact Regression in Simultaneous Slices"

Tubiolo *et al.*

**These supplementary materials include:**

Supplementary Methods

Supplementary Results and Discussion

Supplementary Figures S1 – S18

Supplementary Tables S1 – S7

Supplementary References

### Supplementary Methods

#### Task Procedures

*Self-ordered Working Memory Task (SOT)*

The Self-ordered Working Memory Task (SOT) was modified from previously published work[1,2,3], and consisted of 20 trials, with eight steps of gradually increasing working memory (WM) load in each trial. At the start of a trial, a three-by-three grid of eight simple line drawings of three-dimensional objects was presented, with the center position in the grid left blank. Unique stimuli were used on each of the first 10 trials (80 total stimuli) and were each repeated once in the following 10 trials. Participants were given seven seconds in which to respond on each step. Responses consisted of using an fMRI compatible trackball to position a cursor over one of the objects and select it with a button press. Participants were instructed to select any object on each step that they had not already selected on a previous step (thus, on the first step all possible responses are correct). Once participants made a selection, a white square was displayed around the selected object until a total of nine seconds had elapsed since the beginning of the step, thus ensuring that each step remained the same length regardless of participants' reaction times. At the start of each subsequent step after the first, the objects were pseudo-randomly rearranged in the grid, but with the blank space placed at the location of the previously selected item (thus preventing participants from simply selecting the same location on each step). If participants failed to make a response within seven seconds from the beginning of a step, a white square was displayed around a randomly selected object that would have been a correct response. Participants were instructed to remember this object as if they had selected it themselves, and to continue the trial. If an incorrect selection was made, a red square was displayed over top of the selected object in order to indicate that an error had been made, and the same procedure as in the case of no response was followed. Finally, participants also carried out two trials of a control task, in which one of the objects on each step was marked with an asterisk and participants were instructed to simply select the marked object. In all other respects the display and randomization of stimuli for the control task was identical to the SOT. Unique stimuli were used for each of the control trials, and each trial of either type was preceded by textual instructions indicating whether the upcoming trial was a task trial or a control trial.

#### Variance Decomposition of fMRI Data Subspaces

In addition to performing a subspace decomposition of sICA+FIX denoised data, we sought to characterize the effect of MARSS correction on the distribution of variance between these subspaces. The full description and rationale of this decomposition is described in detail elsewhere (see Supplementary Materials in[4]). Briefly, the decomposition of variance in raw, unprocessed data $Var_{Raw}$ can be written as:

$$Var_{Raw} = \boldsymbol{Var_{Detrend}} + \boldsymbol{Var_{MP24}} + \boldsymbol{Var_{NoiseICA}} + \boldsymbol{Var_{Neural}} + \boldsymbol{Var_{Random}} \quad (1)$$

where $Var_{Detrend}$ is calculated from signal removed via linear detrending, $Var_{MP24}$ is calculated from signal removed by regression of six motion parameters, their squares, derivatives, and squared derivatives, $Var_{NoiseICA}$ is the variance estimated from noise independent components, $Var_{Neural}$ is the variance of the neural signal subspace, and $Var_{Random}$ is the variance of the random noise subspace. With the addition of MARSS correction, a new term $Var_{MARSS}$ is introduced such that $Var_{Raw}$ can now be expressed as

$$Var_{Raw} = \boldsymbol{Var_{Detrend}} + \boldsymbol{Var_{MP24}} + \boldsymbol{Var_{NoiseICA}} + \boldsymbol{Var_{Neural}} + \boldsymbol{Var_{Random}} + \boldsymbol{Var_{MARSS}} \quad (2)$$

Methods for calculating each subspace are described elsewhere (see main text, *2.5.2. Subspace Decomposition of sICA+FIX Denoised Data*). For each resting-state run of each participant in the HCP-Siemens dataset, the variance terms in the above equations were calculated in uncorrected and MARSS-corrected data, the only difference being that $Var_{MARSS}$

cannot be calculated for uncorrected data. Variance for each subspace was calculated as the mean variance of all in-brain voxels.

We were also particularly interested in calculating the percentage of variance explained by the MARSS artifact that was removed from the neural signal subspace as a measure of the magnitude of artifact signal that was incorrectly modeled as true neural signal by sICA+FIX. This was done by calculating the variance of the difference between the uncorrected neural signal subspace and the MARSS-corrected neural signal subspace and dividing by the variance of the MARSS-corrected neural signal subspace.

#### Supplemtary Results and Discussion

##### Variance Decomposition of fMRI Data Subspaces

The results of a variance decomposition of data subspaces in both uncorrected and MARSS-corrected resting-state data from the HCP-Siemens dataset are shown in **Table S7**. Decreases in mean voxel-wise variance were observed in the signal removed by linear detrending, motion parameter regression, and sICA+FIX noise components, while increases in variance were observed in the neural signal subspace and random noise subspace. Standard error of the mean variance slightly decreased in all subspaces. The variance of the difference between the uncorrected and MARSS-corrected neural subspaces was calculated to be 67.857 ± 10.490 (confidence interval reflect 2 * standard error of the mean). Consequently, signal changes in the neural subspace by MARSS correction explains approximately 1.128% of the variance in the neural subspace.

Across all subspaces, the largest decrease in variance was observed in the sICA+FIX noise component subspace. This may indicate that MARSS correction removes a portion of the structured noise that would have been estimated and removed by sICA+FIX noise components. As a result of the reduced signal magnitude being removed by these components, a slightly higher signal variance is observed in the neural signal subspace, likely suggesting the return of true variance to this subspace that had been previously estimated as noise, as well as an increase in the variance of unstructured noise in the random noise subspace.

While these results indicate an overall improvement in both the performance of sICA+FIX and in data quality, this analysis is limited by the fact that the independent component regressors used to performed sICA+FIX denoising were previously estimated on data that had been normalized to a template space (and subsequently applied to unprocessed data in the scanner space). This normalization distorts the simultaneous slice acquisition pattern, and thus sICA+FIX is not expected to be able to identify noise signals following that structure. Additionally, independent component regressors that were estimated on uncorrected data were used on MARSS-corrected data. Therefore, this analysis should be repeated in future work with the added step of estimating sICA+FIX components on MARSS-corrected data, specifically.

#### Supplementary Figures

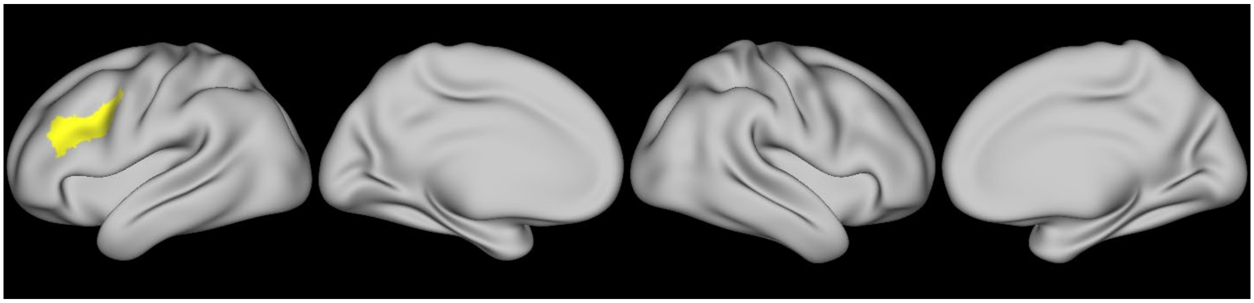

**Figure S1.** Left dorsolateral prefrontal cortex region of interest used for evaluation of changes in group-level t-statistics due to artifact correction.

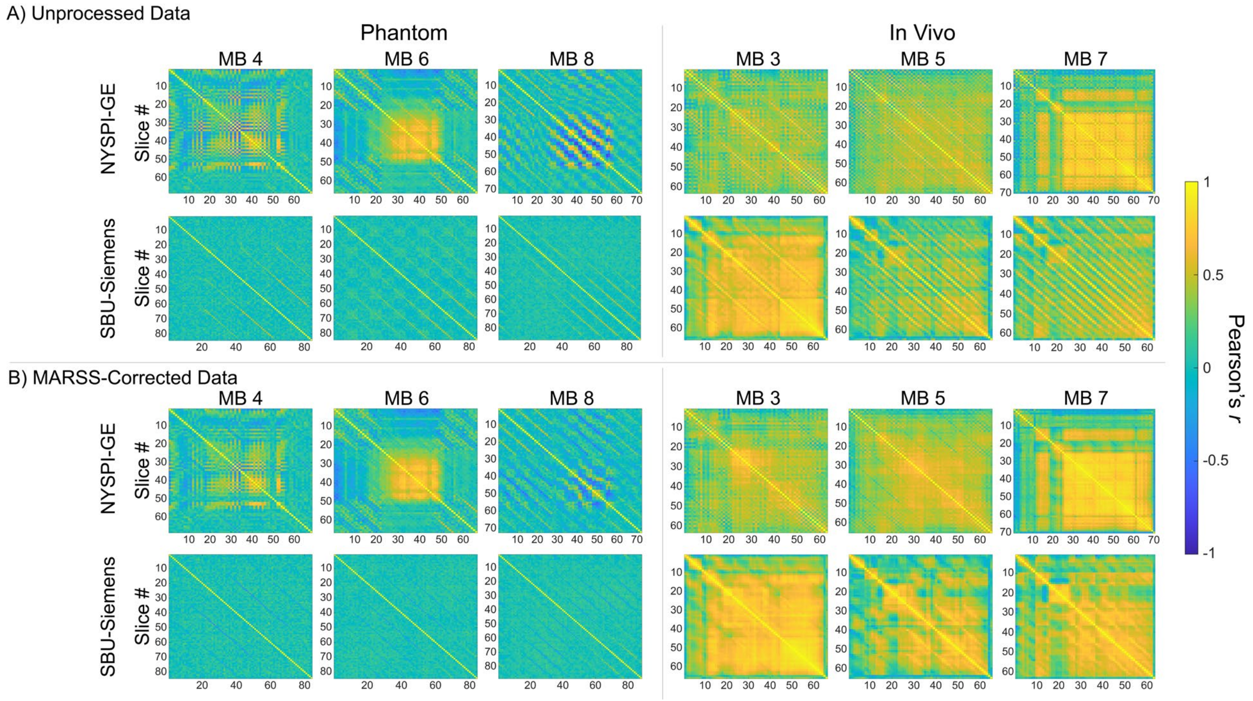


**Figure S2.** Pearson correlation matrices between average signal in all slice pairs in A) unprocessed data and B) MARSS-artifact corrected data, performed in phantom and in vivo at varying multiband (MB) acceleration factors.

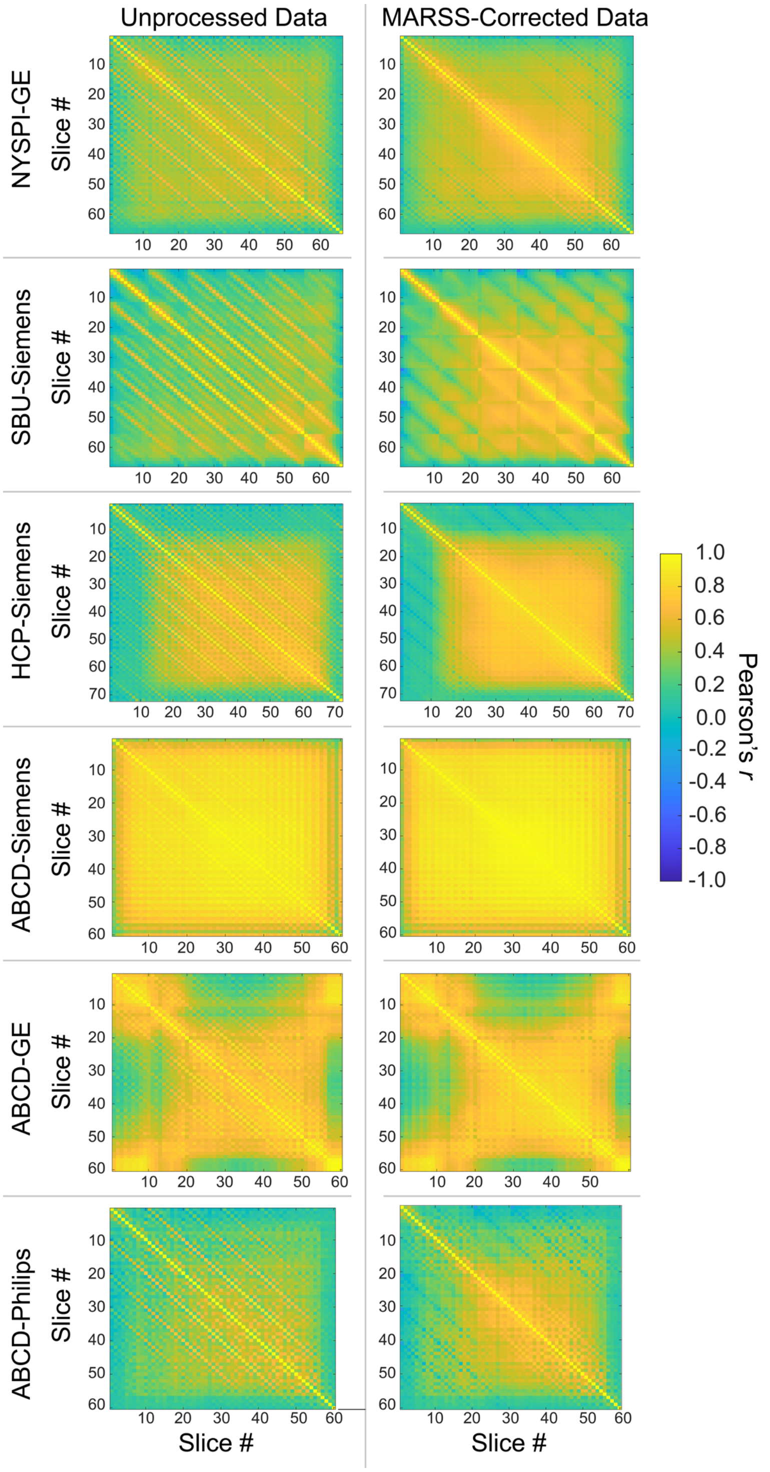


**Figure S3.** Pearson correlation matrices between average signal in all slice pairs in all unprocessed and MARSS-corrected resting-state datasets.

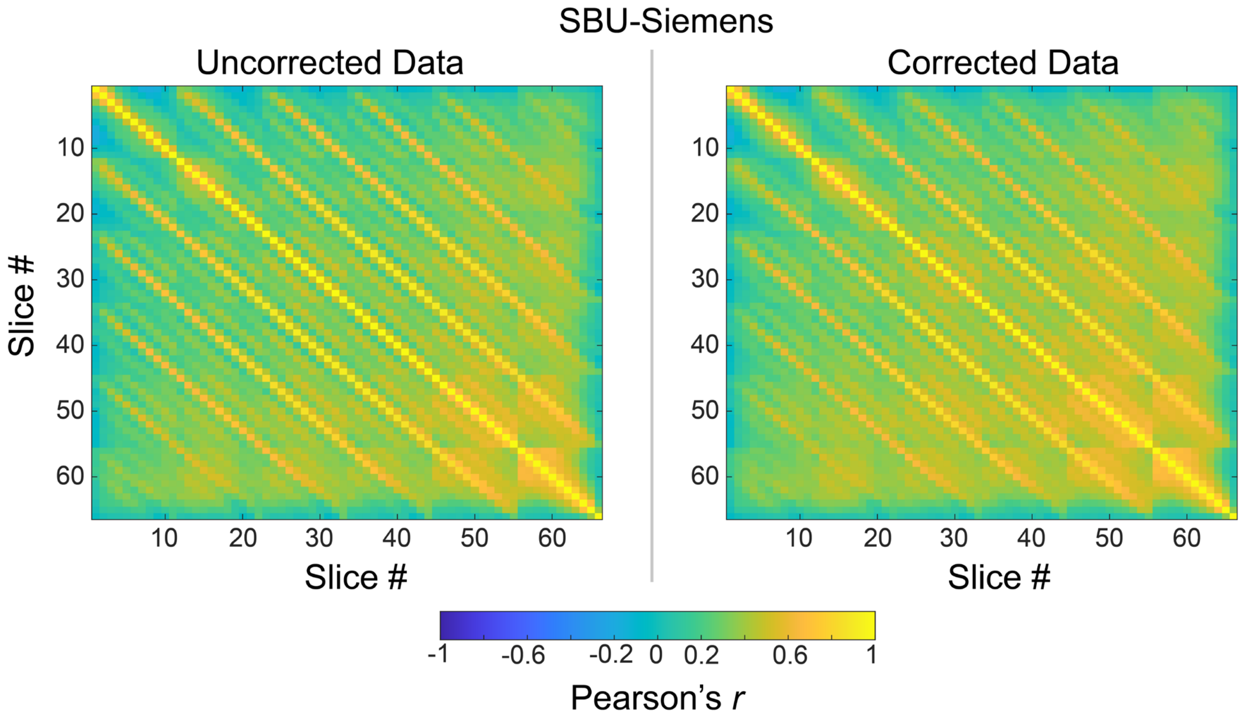


**Figure S4.** Pearson correlation matrices between average signal in all slice pairs in uncorrected data and data corrected with the alternative technique of estimating artifact signal from image background alone. Analysis was performed in resting-state data from SBU-Siemens.

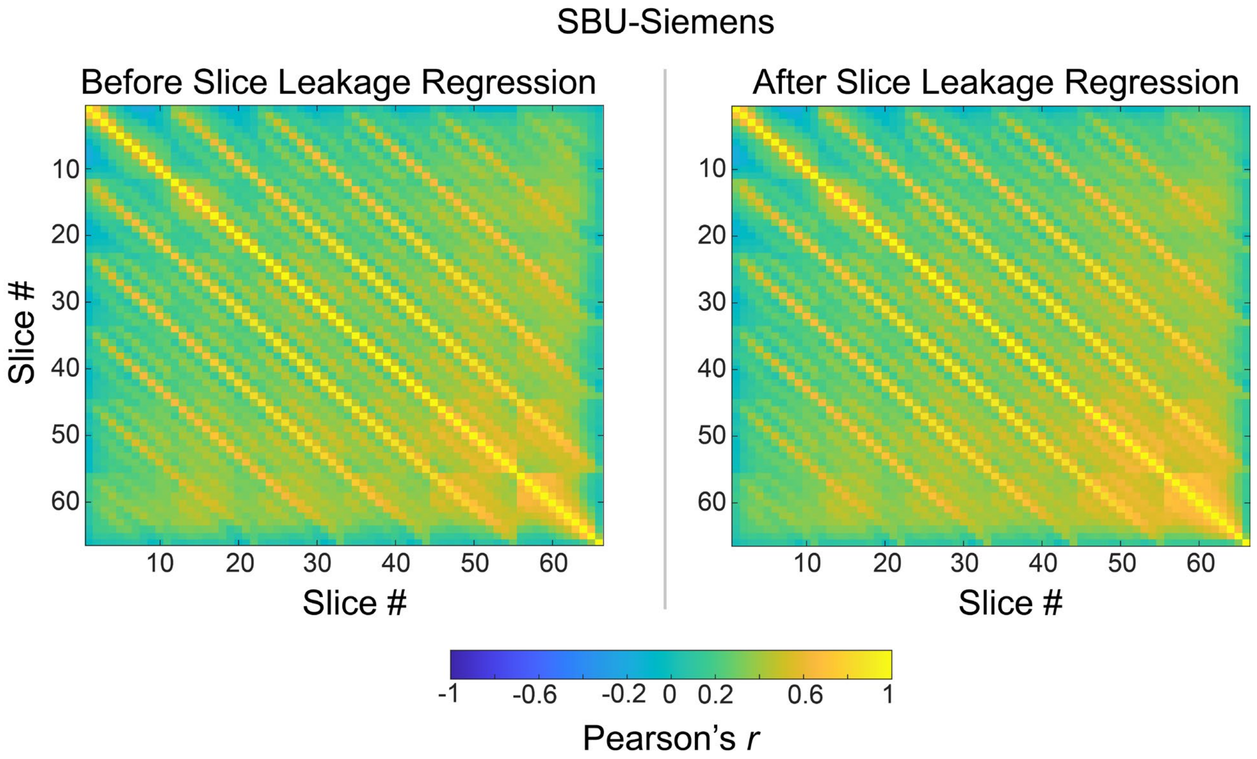


**Figure S5.** Pearson correlation matrices between average signal in all slice pairs in uncorrected data and data corrected by slice leakage regression. Analysis was performed in resting-state data from SBU-Siemens.

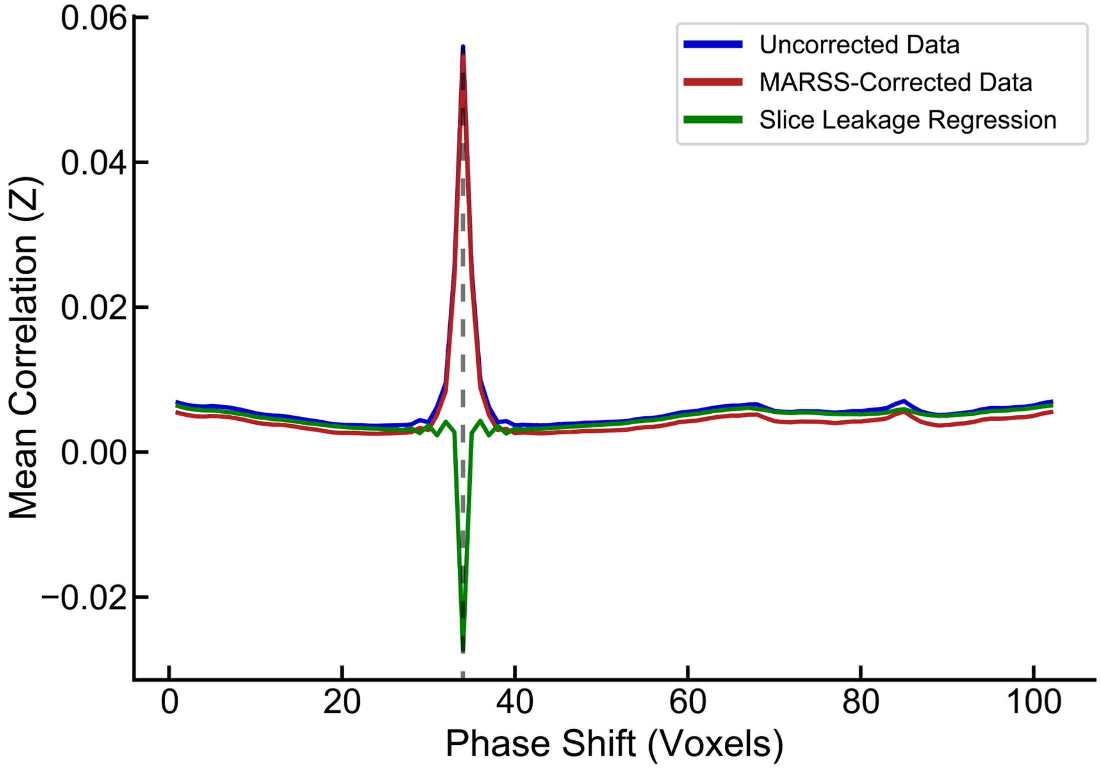


**Figure S6.** Average signal correlations between voxels in simultaneously acquired slices as a function of the phase shift between them during multiband reconstruction in resting-state data from the SBU-Siemens dataset. The phase shift in this dataset was 34 voxels (marked with grey dashed line), meaning that voxels in a given slice overlapped with the voxels that were exactly 34 voxels away along the phase-encode direction during reconstruction. Analysis was performed in uncorrected data (blue line), MARSS-corrected data (red line), and uncorrected data that had undergone slice leakage regression, only (green line).

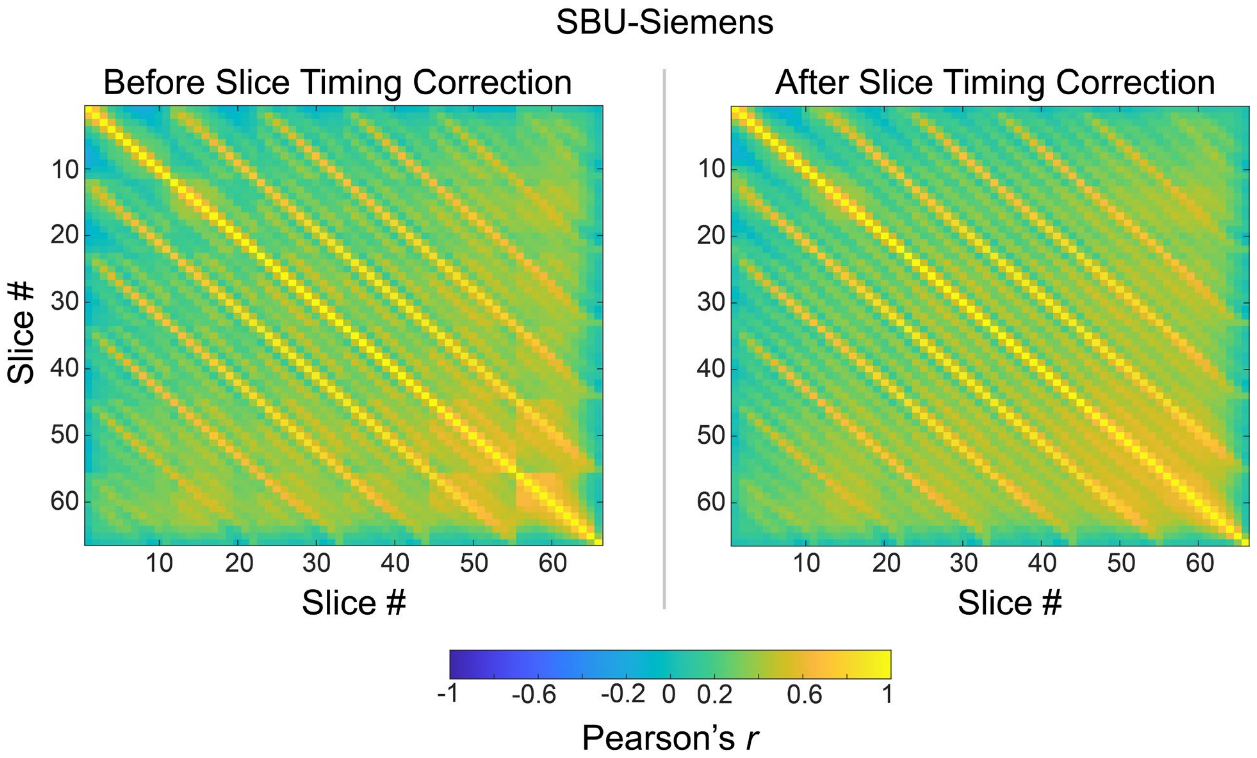


**Figure S7.** Pearson correlation matrices between average signal in all slice pairs in uncorrected data and data that has undergone slice timing correction. Analysis was performed in resting-state data from SBU-Siemens.

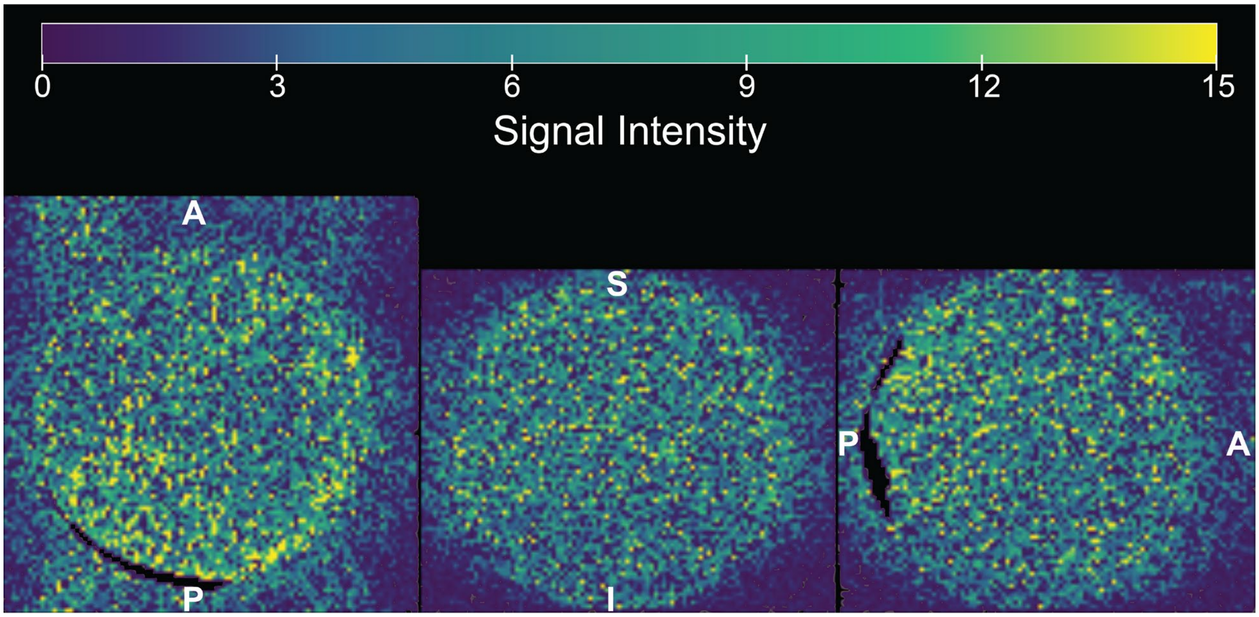


**Figure S8.** Average spatial distribution of the isolated artifact signal in SBU-Siemens phantom data collected with a multiband factor of 6. The black portion of the image occurred to an "amplifier blocking" effect in which signal intensity reached the maximum value in those voxels, resulting on a variance of 0, preventing the artifact from being estimated in those voxels.

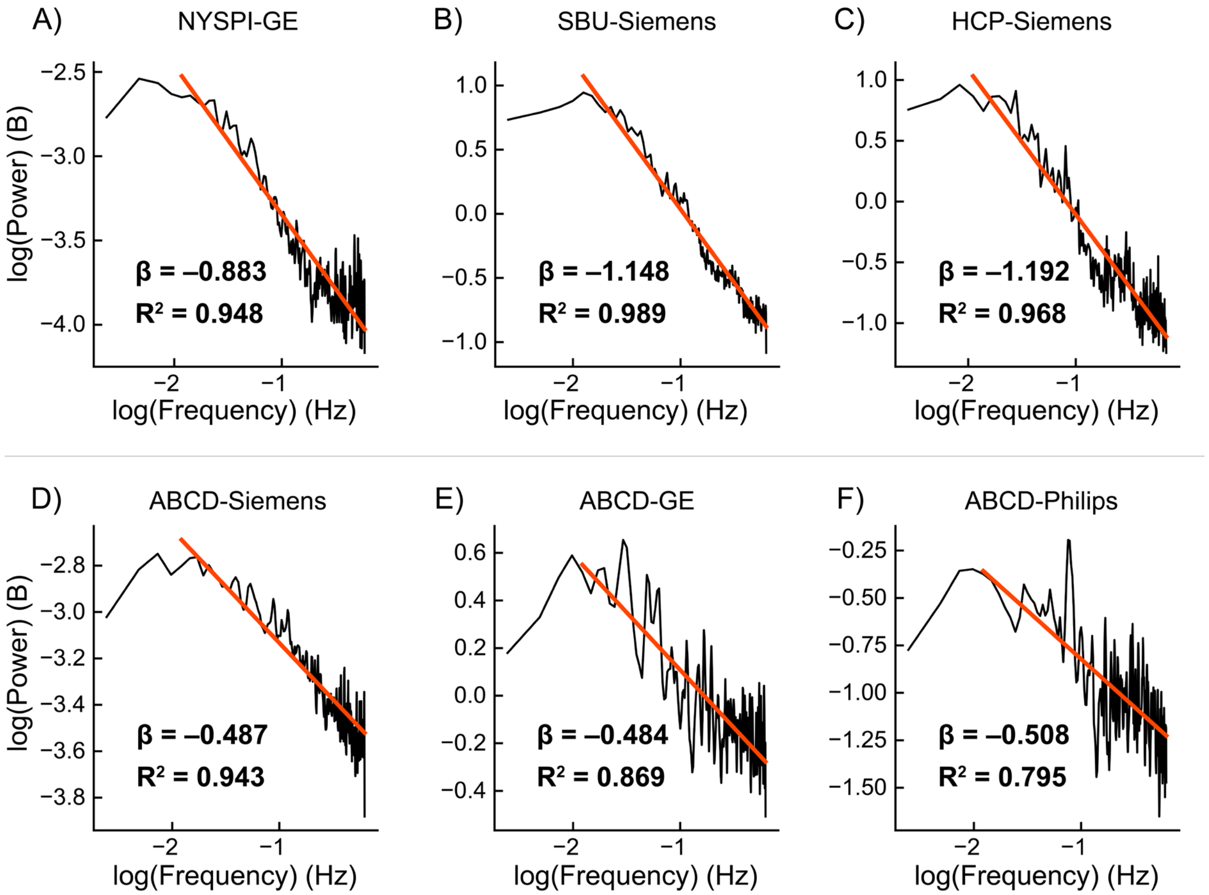


**Figure S9.** Log-log transformed power spectral density of the isolated artifact timeseries concatenated across all participants in resting-state fMRI data from A) NYSPI-GE, B) SBU-Siemens, C) HCP-Siemens, D) ABCD-Siemens, E) ABCD-GE, and F) ABCD-Philips. Plot annotation denotes the slope (β) and $R^2$ of each linear fit (red line).

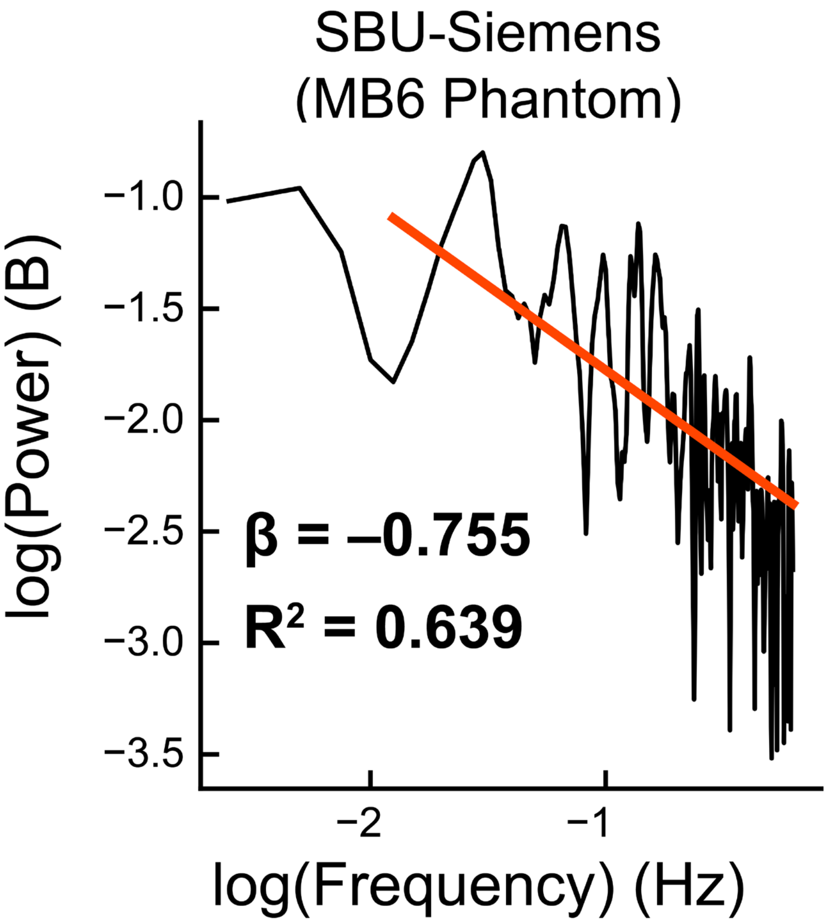


**Figure S10.** Log-log transformed power spectral density of the isolated artifact timeseries in SBU-Siemens phantom data collected with a multiband factor of 6. Plot annotation denotes the slope (β) and $R^2$ of the linear fit (red line).

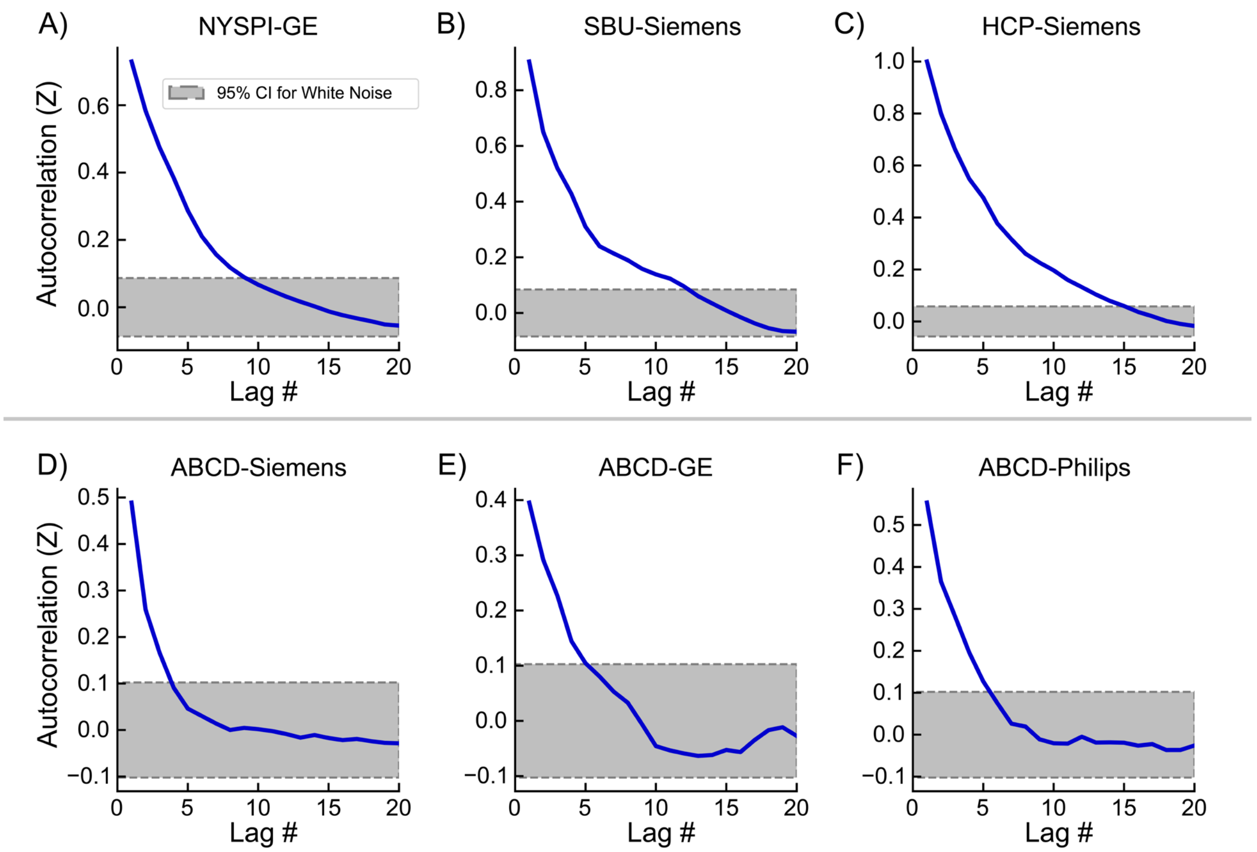


**Figure S11.** Mean, Z-transformed autocorrelation of the isolated artifact timeseries as a function of lag in resting-state fMRI data from A) NYSPI-GE, B) SBU-Siemens, C) HCP-Siemens, D) ABCD-Siemens, E) ABCD-GE, and F) ABCD-Philips. Grey area with dotted borders denotes the 95% confidence interval for significant autocorrelation over and above that of white noise.

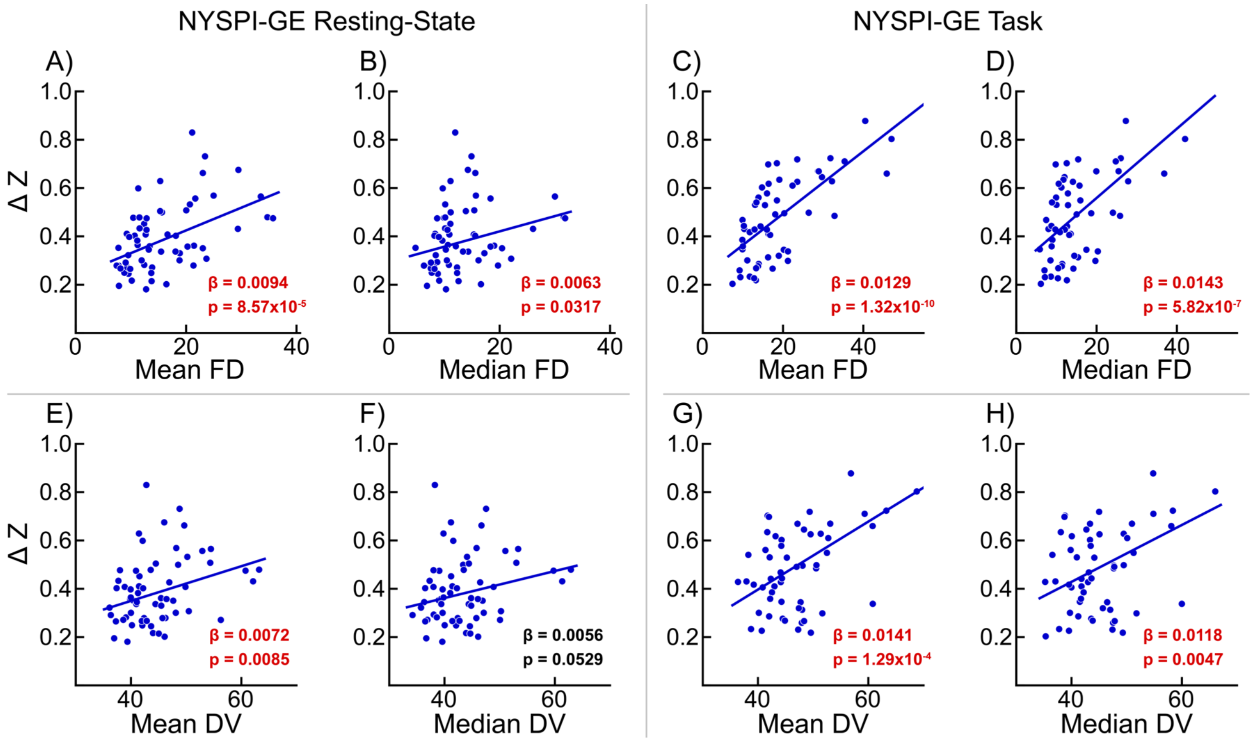


**Figure S12.** Associations between artifact magnitude, measured as the difference in Z-transformed Pearson correlation between simultaneously acquired slices and adjacent slices, and mean framewise displacement (FD; A, C), median FD (B, D), mean DV (E, G), and median DV (F, H). Inlaid text denotes slope (β) and p-value (p) of linear fit, with red text denoting statistical significance at $p<0.05$. Analyses were performed in the NYSPI-GE resting-state and task-based datasets.

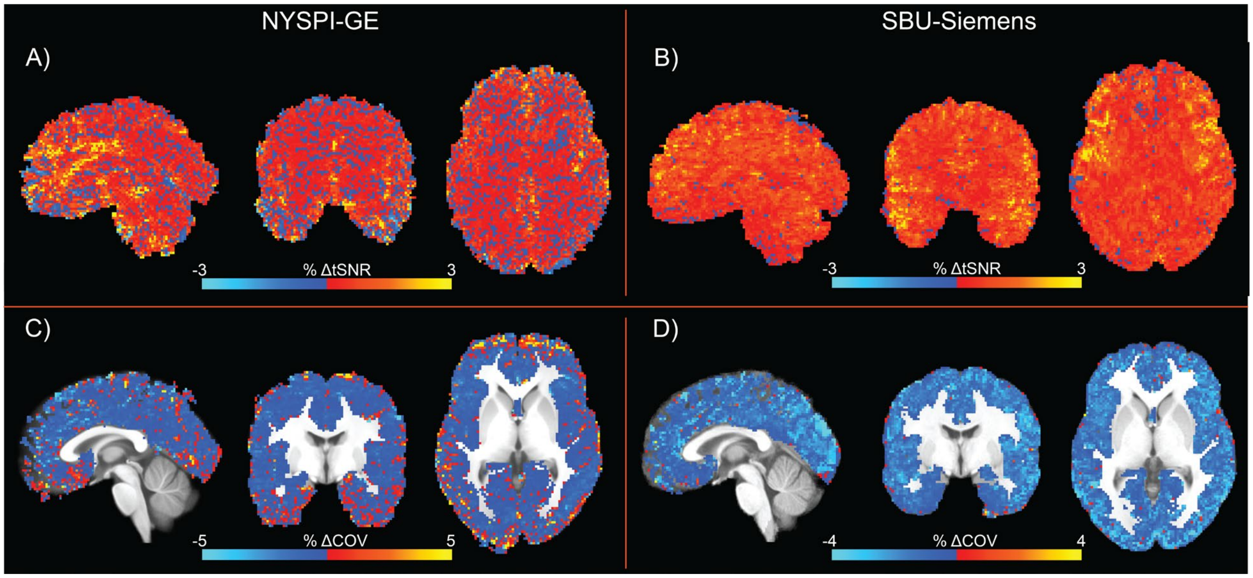


**Figure S13**. Mean, voxel-wise percent change in temporal signal-to-noise ratio (tSNR; A,B) and Coefficient of Variation (COV, C,D) as a result of MARSS artifact correction in preprocessed, task-based fMRI data from NYSPI-GE (A,C) and SBU-Siemens (B,D).

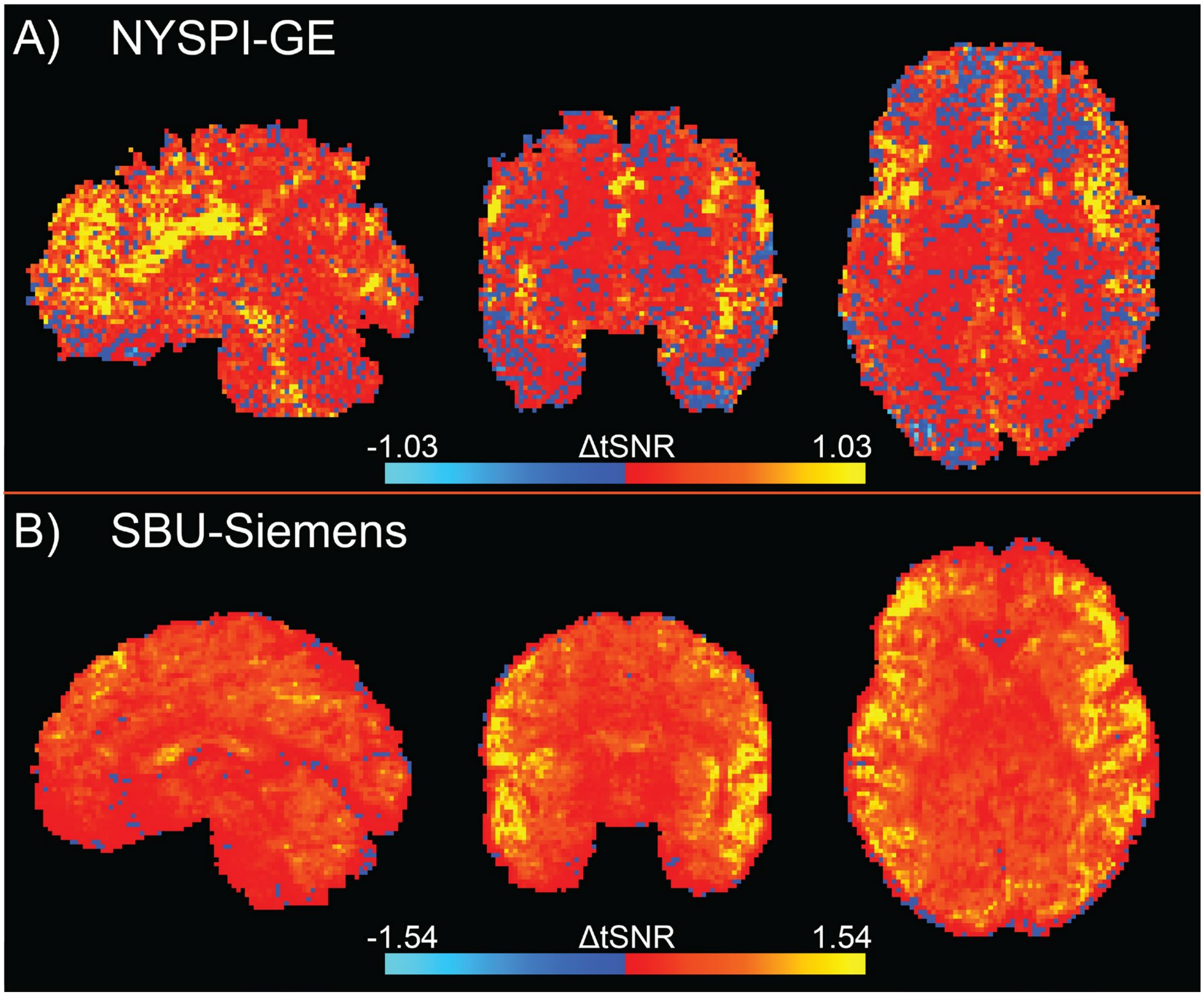


**Figure S14.** Mean, voxel-wise change in temporal signal-to-noise ratio as a result of MARSS artifact correction in preprocessed, resting-state fMRI data from A) NYSPI-GE and B) SBU-Siemens.

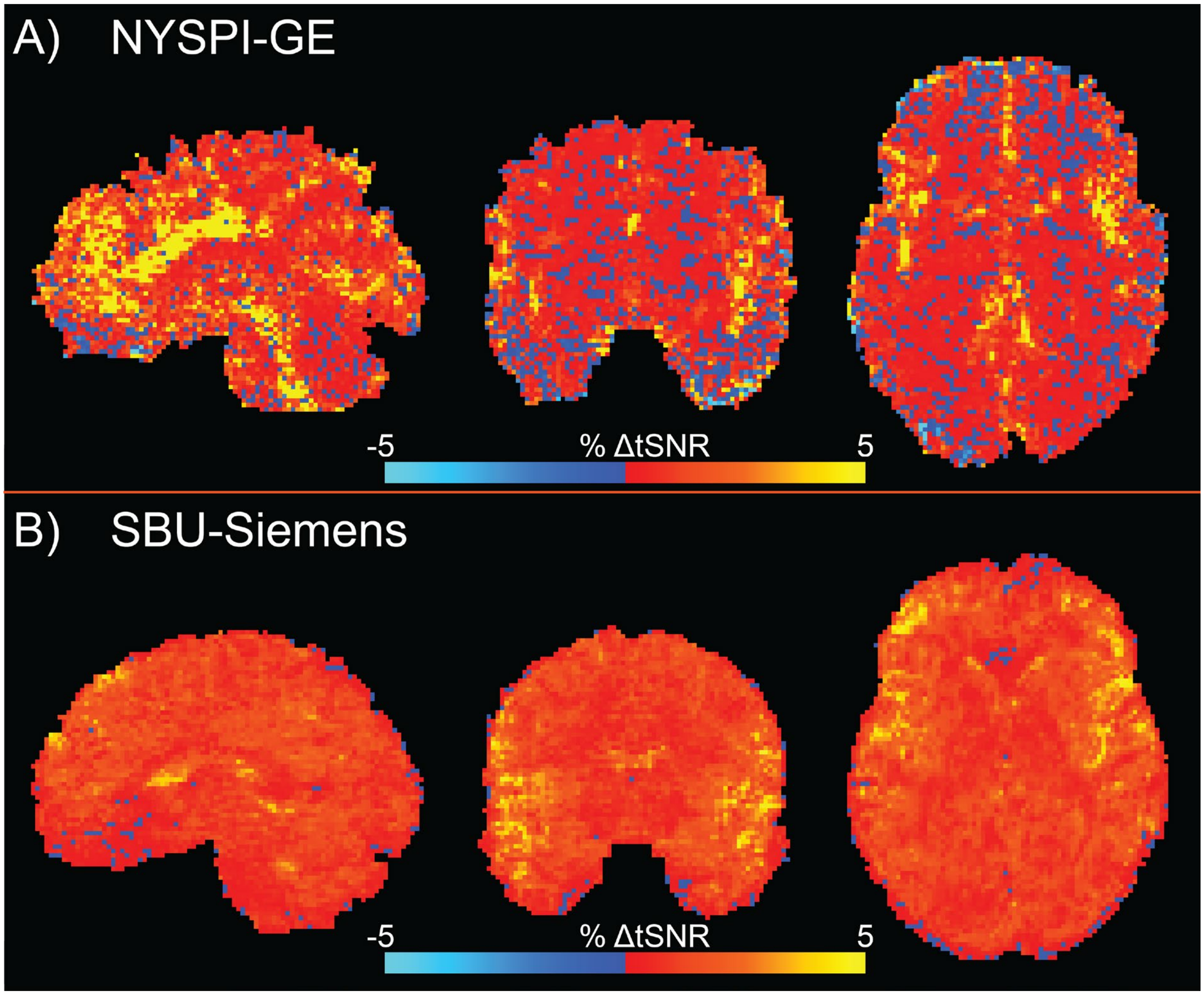


**Figure S15.** Mean, voxel-wise percent change in temporal signal-to-noise ratio as a result of MARSS artifact correction in preprocessed, resting-state fMRI data from A) NYSPI-GE and B) SBU-Siemens.

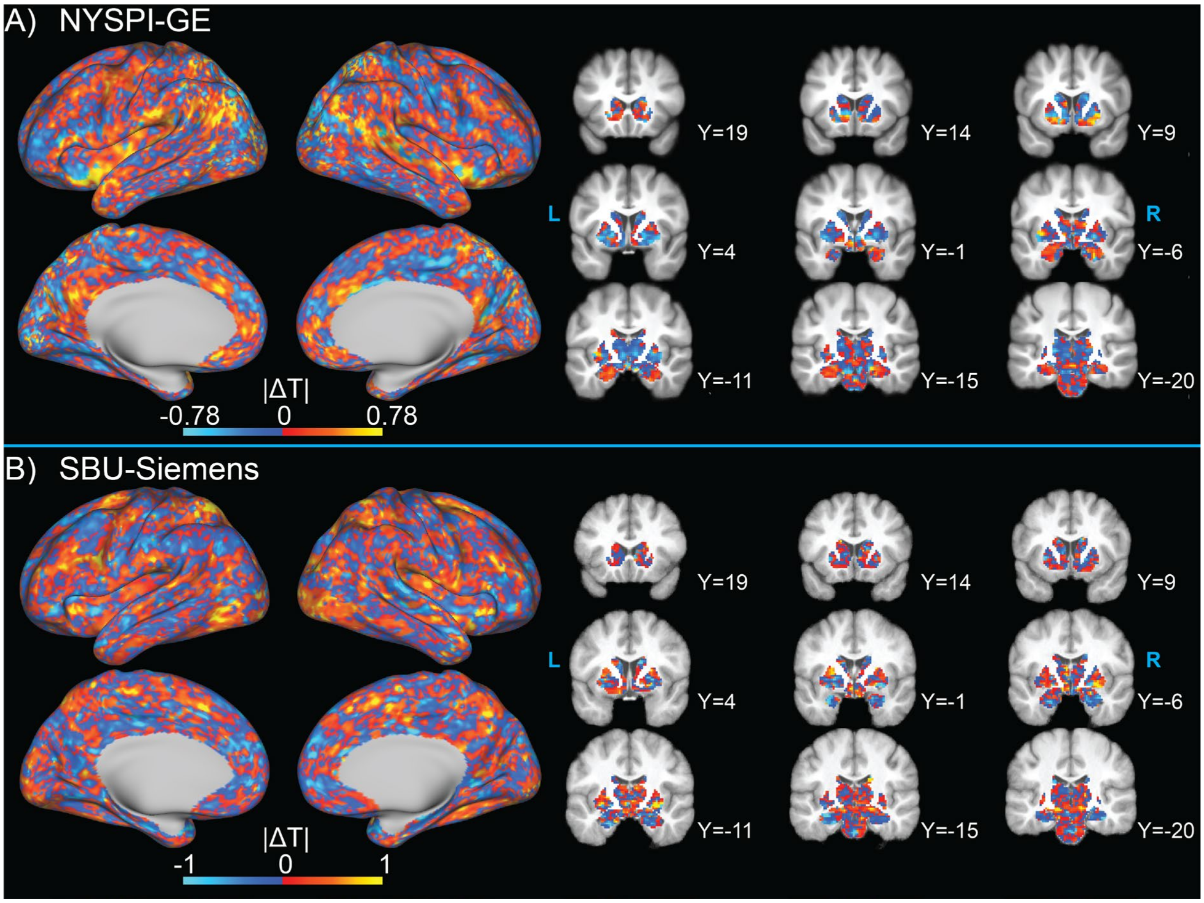


**Figure S16.** Change in t-statistic magnitude in between-participants analysis of task-evoked activation as a result of MARSS artifact correction in data from A) NYSPI-GE and B) SBU-Siemens.

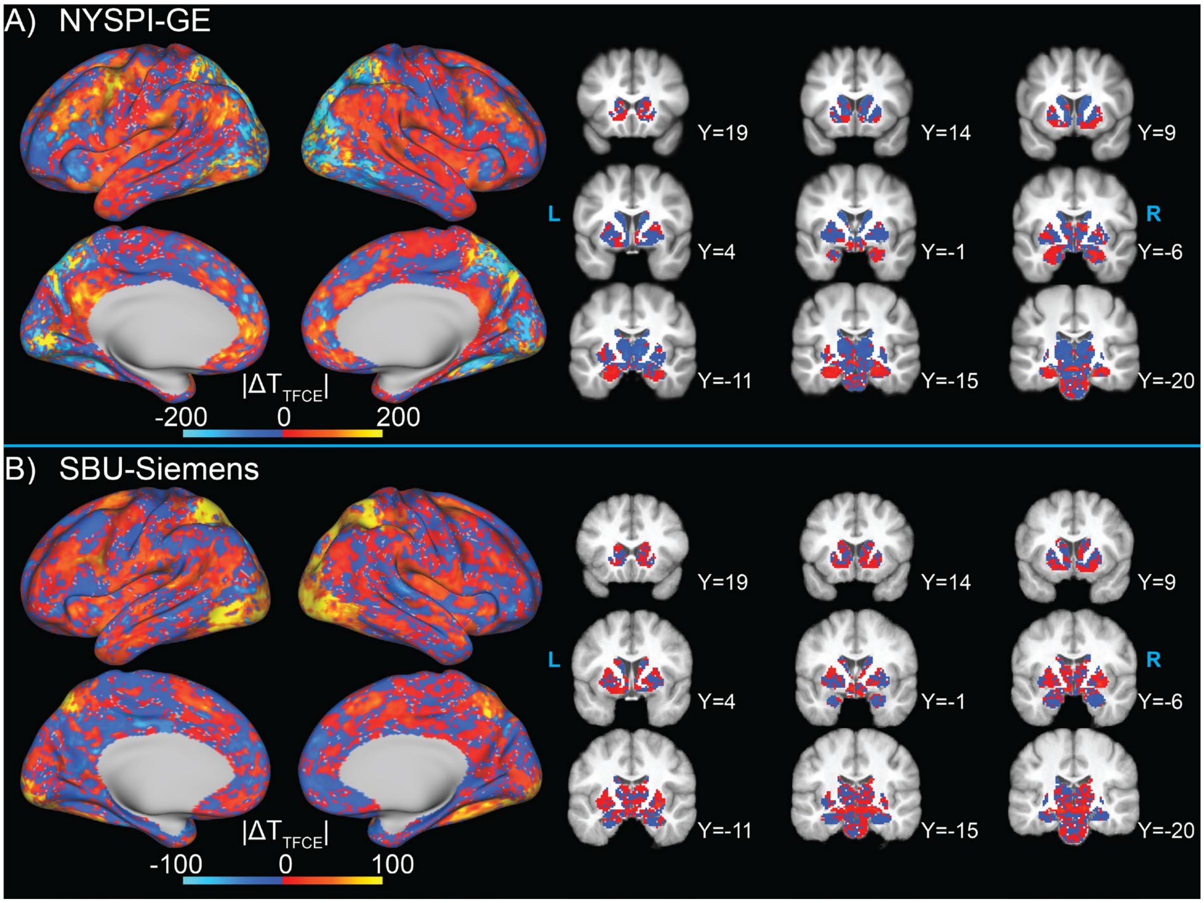


**Figure S17.** Change in t-statistic magnitude in between-participants analysis of task-evoked activation as a result of MARSS artifact correction in data collected from A) NYSPI-GE and B) SBU-Siemens. T-statistic maps before and after correction were generated using threshold-free cluster enhancement (TFCE) during permutation testing.

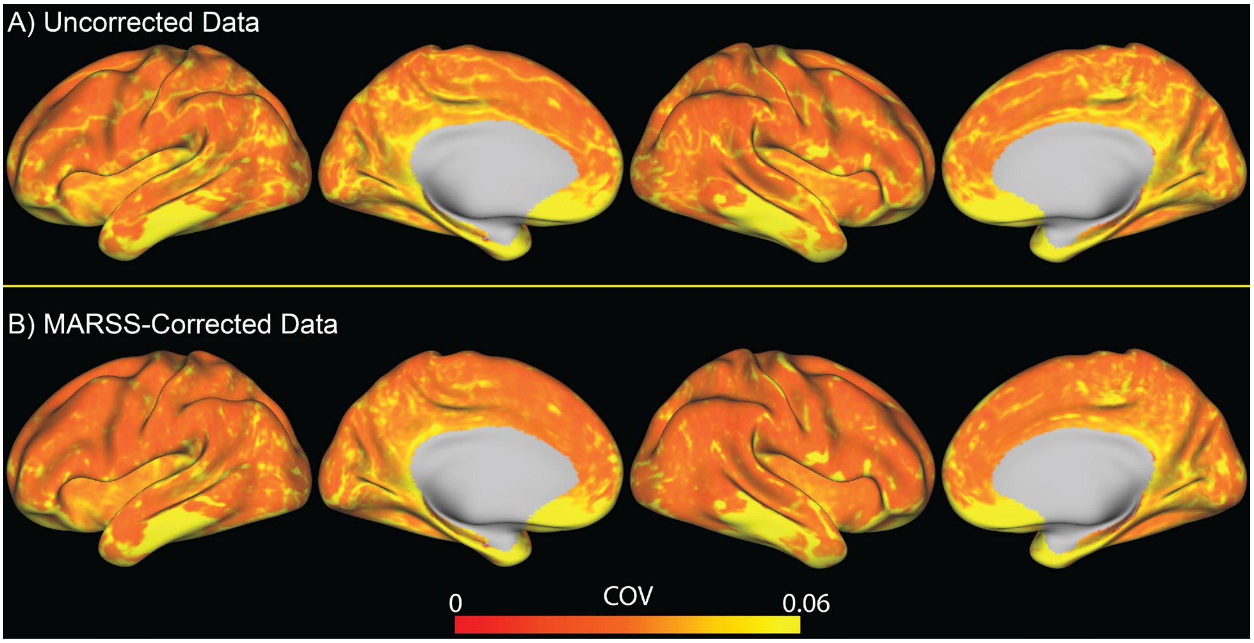


**Figure S18.** Cortical coefficient of variance (COV) before and after MARSS artifact correction in a task-based fMRI run from a single participant in the NYSPI-GE dataset Clear stripes of high COV in uncorrected data resulted in the exclusion of these voxels from this participant's volume-to-surface mapping process.

#### Supplementary Tables

**Table S1.** Demographic information for all datasets used in this study.

| Dataset | N | Mean Age [SD] | M/F | Race | Ethnicity |
|---|---|---|---|---|---|
| SBU | 10 | 28 [9.43] | 10/0 | 7W/1AA/0AS/2 Other | 2H/8NH |
| NYSPI | 66 | 32 [11.13] | 48/18 | 17W/31AA/6AS/4MR/8 Other/Unknown | 17H/49NH |
| HCP | 25 | 29.08 [3.73] | 18/7 | 18W/6AA/1AS | 24NH/1 Unknown |
| ABCD | 35 | 10.29 [1.16] | 18/17 | 20W/5AA/2AS/1MR/1 Other/3 Unknown | 12H/20NH/3 Unknown |

SBU = Stony Brook University; NYSPI = New York State Psychiatric Institute; HCP = Human Connectome Project; ABCD = Adolescent Brain Cognitive Development; SD = Standard Deviation; M/F = Male/Female; W = White; AA = African American; AS = Asian; MR = Multiracial; H = Hispanic; NH = Non-Hispanic

**Note:** for three ABCD participants, only information on sex was available and was used in place of gender. Race/ethnicity information was missing for these participants as well, and they are reported "Unknown."

**Table S2**. Mean Pearson correlations between simultaneously acquired (SA+) slices and adjacent-to-simultaneous (Adj) slices in uncorrected and MARSS-corrected data for all resting-state and task-based datasets.

| | Uncorrected Data | | | MARSS-Corrected Data | | |
|---|---|---|---|---|---|---|
| **Dataset** | $r_{SA+}$ | $r_{Adj}$ | $\Delta r_{(SA+\,-\,Adj)}$ $(\Delta Z)$ | $r_{SA+}$ | $r_{Adj}$ | $\Delta r_{(SA+\,-\,Adj)}$ $(\Delta Z)$ |
| NYSPI-GE RS | 0.608 | 0.228 | 0.380 (0.400) | 0.379 | 0.447 | -0.068 (-0.068) |
| NYSPI-GE SOT | 0.604 | 0.146 | 0.458 (0.494) | 0.293 | 0.378 | -0.085 (-0.085) |
| SBU-Siemens RS | 0.682 | 0.283 | 0.399 (0.423) | 0.395 | 0.433 | -0.038 (-0.038) |
| SBU-Siemens SOT | 0.667 | 0.234 | 0.433 (0.464) | 0.256 | 0.299 | -0.043 (-0.043) |
| HCP-Siemens RS | 0.631 | 0.217 | 0.414 (0.440) | 0.429 | 0.465 | -0.036 (-0.036) |
| HCP-Siemens WM | 0.574 | 0.109 | 0.465 (0.504) | 0.261 | 0.311 | -0.050 (-0.050) |
| ABCD-Siemens RS | 0.870 | 0.600 | 0.270 (0.276) | 0.836 | 0.861 | -0.025 (-0.025) |
| ABCD-GE RS | 0.667 | 0.438 | 0.229 (0.233) | 0.599 | 0.578 | 0.021 (0.021) |
| ABCD-Philips RS | 0.540 | 0.168 | 0.372 (0.391) | 0.288 | 0.035 | -0.062 (-0.062) |

RS = Resting-State; SOT = Self-Ordered Working Memory Task; WM = Working Memory Task; $r_{SA+}$ = Mean Pearson correlation between simultaneously acquired slices; $r_{Adj}$ = Mean Pearson correlation between adjacent slices; $\Delta r_{(SA+\,-\,Adj)}$ = Mean difference in Pearson correlation between simultaneously acquired slices and adjacent slices; $\boldsymbol{\Delta Z}$ = Mean difference in Z-transformed Pearson correlation between simultaneously acquired slices and adjacent slices.

**Table S3.** Mean Pearson correlations between simultaneously acquired (SA+) slices and adjacent-to-simultaneous (Adj) slices in phantom and in vivo resting-state data with varying multiband (MB) factors in uncorrected and MARSS-corrected data.

| Dataset | Phantom /In Vivo | MB Factor | Uncorrected Data $r_{SA+}$ | Uncorrected Data $r_{Adj}$ | Uncorrected Data $\Delta r_{(SA+\,-\,Adj)}$ $(\Delta Z)$ | MARSS-Corrected Data $r_{SA+}$ | MARSS-Corrected Data $r_{Adj}$ | MARSS-Corrected Data $\Delta r_{(SA+\,-\,Adj)}$ $(\Delta Z)$ |
|---|---|---|---|---|---|---|---|---|
| NYSPI-GE | In Vivo | 3 | 0.539 | 0.249 | 0.290 (0.298) | 0.245 | 0.368 | -0.123 (-0.124) |
| | | 5 | 0.569 | 0.188 | 0.364 (0.381) | 0.324 | 0.428 | -0.104 (-0.104) |
| | | 7 | 0.592 | 0.335 | 0.257 (0.263) | 0.467 | 0.482 | -0.015 (-0.015) |
| | Phantom | 4 | 0.214 | 0.009 | 0.205 (0.208) | 0.021 | 0.067 | -0.046 (-0.046) |
| | | 6 | 0.226 | 0.106 | 0.120 (0.121) | 0.037 | 0.106 | -0.069 (-0.069) |
| | | 8 | 0.215 | 0.068 | 0.147 (0.148) | -0.094 | -0.004 | -0.090 (-0.090) |
| SBU-Siemens | In Vivo | 3 | 0.689 | 0.448 | 0.241 (0.246) | 0.501 | 0.523 | -0.022 (-0.022) |
| | | 5 | 0.624 | 0.225 | 0.399 (0.422) | 0.321 | 0.334 | -0.013 (-0.013) |
| | | 7 | 0.692 | 0.282 | 0.410 (0.436) | 0.435 | 0.459 | -0.024 (-0.024) |
| | Phantom | 4 | 0.212 | 0.007 | 0.205 (0.208) | -0.112 | 0.035 | -0.147 (-0.148) |
| | | 6 | 0.213 | 0.057 | 0.156 (0.158) | -0.056 | 0.049 | -0.105 (-0.106) |
| | | 8 | 0.189 | 0.044 | 0.145 (0.146) | -0.020 | 0.052 | -0.072 (-0.072) |

$r_{SA+}$ = Mean Pearson correlation between simultaneously acquired slices; $r_{Adj}$ = Mean Pearson correlation between adjacent slices; $\Delta r_{(SA+\,-\,Adj)}$ = Mean difference in Pearson correlation between simultaneously acquired slices and adjacent slices; $\Delta Z$ = Mean difference in Z-transformed Pearson correlation between simultaneously acquired slices and adjacent slices.

**Table S4**. Results of Wilcoxon Signed Rank tests to determine the significance of the reduction in correlation difference between simultaneously acquired slices and adjacent slices following MARSS artifact correction.

| Dataset | n | W | p |
|---|---|---|---|
| NYSPI-GE RS | 232 | 27,028 | $8.21 \times 10^{-40}$ |
| NYSPI-GE SOT | 186 | 17,578 | $1.94 \times 10^{-32}$ |
| SBU-Siemens RS | 42 | 903 | $1.65 \times 10^{-8}$ |
| SBU-Siemens SOT | 39 | 780 | $5.26 \times 10^{-8}$ |
| HCP-Siemens RS | 100 | 5,050 | $3.90 \times 10^{-18}$ |
| HCP-Siemens WM | 50 | 1,275 | $7.56 \times 10^{-10}$ |
| ABCD-Siemens RS | 81 | 3,321 | $5.36 \times 10^{-15}$ |
| ABCD-GE RS | 9 | 45 | 0.0039 |
| ABCD-Philips RS | 21 | 231 | $5.96 \times 10^{-5}$ |

RS = Resting-State; SOT = Self-Ordered Working Memory Task; WM = Working Memory Task; n = number of runs in each dataset; W = Wilcoxon signed rank test statistic; p = p-value

**Table S5.** Change in temporal signal-to-noise ratio (tSNR) as a result of MARSS artifact correction in all unprocessed, resting-state and task-based fMRI datasets. Elevations in tSNR were tested for statistical significance via Wilcoxon Signed Rank tests.

| Dataset | $tSNR_{uncorrected}$ | $tSNR_{MARSS}$ | $\Delta tSNR$ | n | W | p |
|---|---|---|---|---|---|---|
| NYSPI-GE RS | 18.970 | 19.027 | 0.057 | 232 | 27,028 | $8.201 \times 10^{-40}$ |
| NYSPI-GE SOT | 17.649 | 17.705 | 0.056 | 186 | 17,578 | $1.944 \times 10^{-32}$ |
| SBU-Siemens RS | 29.161 | 29.363 | 0.202 | 42 | 903 | $1.648 \times 10^{-8}$ |
| SBU-Siemens SOT | 27.192 | 27.416 | 0.224 | 39 | 780 | $5.255 \times 10^{-8}$ |
| HCP-Siemens RS | 16.614 | 16.665 | 0.051 | 100 | 5,050 | $3.897 \times 10^{-18}$ |
| HCP-Siemens WM | 19.456 | 19.535 | 0.079 | 50 | 1,275 | $7.557 \times 10^{-10}$ |
| ABCD-Siemens RS | 23.921 | 24.090 | 0.169 | 81 | 3,321 | $5.363 \times 10^{-15}$ |
| ABCD-GE RS | 25.336 | 25.210 | 0.126 | 9 | 45 | 0.004 |
| ABCD-Philips RS | 23.822 | 24.043 | 0.221 | 21 | 231 | $5.957 \times 10^{-5}$ |

RS = Resting-State; SOT = Self-Ordered Working Memory Task; WM = Working Memory Task; $tSNR$ = temporal signal-to-noise ratio; n = number of runs in each dataset; W = Wilcoxon Signed Rank test statistic; p = p-value.

**Table S6**. Mean Pearson correlations between simultaneously acquired (SA+) slices and adjacent-to-simultaneous (Adj) slices in subspaces of sICA+FIX denoised, resting-state data from the HCP-Siemens dataset.

| | **Uncorrected Data** | | | **MARSS-Corrected Data** | | | **Pre-MARSS – Post-MARSS** | | |
|---|---|---|---|---|---|---|---|---|---|
| **Data** | $r_{SA+}$ | $r_{Adj}$ | $\Delta r_{(SA+\,-\,Adj)}$ $(\Delta Z)$ | $r_{SA+}$ | $r_{Adj}$ | $\Delta r_{(SA+\,-\,Adj)}$ $(\Delta Z)$ | $r_{SA+}$ | $r_{Adj}$ | $\Delta r_{(SA+\,-\,Adj)}$ $(\Delta Z)$ |
| sICA+FIX Cleaned | 0.517 | 0.341 | 0.176 (0.178) | 0.396 | 0.398 | -0.002 (-0.002) | 0.960 | -0.017 | 0.977 (2.232) |
| Neural Signal Subspace | 0.707 | 0.626 | 0.081 (0.081) | 0.622 | 0.624 | -0.002 (-0.002) | 0.932 | -0.016 | 0.948 (1.806) |
| Random Noise Subspace | 0.275 | -0.020 | 0.295 (0.304) | 0.052 | 0.053 | -0.001 (-0.001) | 0.967 | -0.016 | 0.983 (2.382) |
| sICA+FIX Noise Components | 0.634 | 0.120 | 0.514 (0.568) | 0.450 | 0.428 | 0.022 (0.022) | 0.989 | -0.007 | 0.996 (3.061) |
| Neural Subspace + GSR | 0.052 | -0.109 | 0.161 (0.162) | -0.080 | -0.076 | -0.004 (-0.004) | 0.914 | -0.029 | 0.943 (1.767) |

$r_{SA+}$ = Mean Pearson correlation between simultaneously acquired slices; $r_{Adj}$ = Mean Pearson correlation between adjacent slices; $\Delta r_{(SA+\,-\,Adj)}$ = Mean difference in Pearson correlation between simultaneously acquired slices and adjacent slices; $\boldsymbol{\Delta Z}$ = Mean difference in Z-transformed Pearson correlation between simultaneously acquired slices and adjacent slices.

**Table S7**. Decomposition of mean voxel-wise variance across participants in data subspaces, performed in both uncorrected and MARSS corrected resting-state data in the HCP-Siemens dataset.

| | **Subspace Variance (Mean ± 2*Standard Error)** | | | | | |
|---|---|---|---|---|---|---|
| | $Var_{Detrend}$ | $Var_{MP24}$ | $Var_{NoiseICA}$ | $Var_{Neural}$ | $Var_{Random}$ | $Var_{MARSS}$ |
| **Uncorrected Data** | 124,206.114 ± 2,857.174 | 96,332.605 ± 1,674.362 | 27,583.954 ± 359.005 | 6,006.277 ± 80.249 | 61,903.802 ± 638.074 | N/A |
| **MARSS-Corrected Data** | 124,186.625 ± 2,857.116 | 96,323.032 ± 1,674.039 | 25,968.731 ± 349.502 | 6,016.725 ± 80.099 | 62,077.783 ± 640.134 | 1,522.526 ± 22.692 |

$Var_{Detrend}$ = Variance calculated on signal removed via linear detrending; $Var_{MP24}$ = Variance calculated on signal removed via regression of 24 motion parameters; $Var_{NoiseICA}$ = Variance calculated on sICA+FIX noise components; $Var_{Neural}$ = Variance of the neural signal subspace; $Var_{Random}$ = Variance of the random noise subspace; $Var_{MARSS}$ = Variance of the signal removed via MARSS correction on raw, unprocessed data.

#### Supplementary References


1. Van Snellenberg JX, Conway AR, Spicer J, Read C, Smith EE. Capacity estimates in working memory: Reliability and interrelationships among tasks. *Cogn Affect Behav Neurosci* **14**, 106-116 (2014).

2. Van Snellenberg JX*, et al.* Dynamic shifts in brain network activation during supracapacity working memory task performance. *Hum Brain Mapp* **36**, 1245-1264 (2015).

3. Van Snellenberg JX*, et al.* Mechanisms of Working Memory Impairment in Schizophrenia. *Biol Psychiatry* **80**, 617-626 (2016).

4. Glasser MF*, et al.* Using temporal ICA to selectively remove global noise while preserving global signal in functional MRI data. *NeuroImage* **181**, 692-717 (2018).
